# Charged mutations in the FUS low-complexity domain modulate condensate ageing kinetics

**DOI:** 10.1101/2025.03.26.645197

**Authors:** Eduardo Pedraza, Daniel Hoyos, Alejandro Feito, Francisco Gámez, Ignacio Sanchez-Burgos, Rosana Collepardo-Guevara, Andrés R. Tejedor, Jorge R. Espinosa

## Abstract

The assembly of biomolecular condensates is tightly regulated by the intracellular environment. Disruptions in the balance between condensate formation and dissolution—such as irreversible aggregation of low-complexity aromatic-rich kinked segments (LARKS)—have been implicated in multiple neuropathologies. Here, we employ non-equilibrium, residue-resolution coarse-grained simulations to investigate how specific mutations in FUS, an RNA-binding protein associated with amyotrophic lateral sclerosis and frontotemporal dementia, modulate its phase separation propensity and transition into insoluble aggregates. Our simulations reveal that mutations increasing the content of negatively charged amino acids in the low-complexity domain slow down inter-protein *β*-sheet accumulation, while preserving the phase diagram and viscoelastic properties of the wild-type sequence. Conversely, mutations increasing the arginine content accelerate disorder-to-order LARKS transitions, driving rapid formation of amorphous kinetically trapped aggregates. Our computational approach thus provides molecular-level insights into how specific amino acid mutations and associated intermolecular interactions control the ageing kinetics of protein condensates, promoting aberrant solid-like phases.

---

Intracellular compartmentalisation is achieved through the intricate interplay of membrane-bound and membraneless organelles, being the latter also known as biomolecular condensates^1–3^. The formation of biomolecular condensates in cells is thought to occur via liquidliquid phase separation (LLPS) of proteins and nucleic acid solutions. The thermodynamic stability and material properties of these condensates are controlled by the balance between homotypic and heterotypic interactions among proteins, nucleic acids, ions, and water^1,4,5^. The continuous assembly and disassembly of biomolecular condensates has been suggested to allow the high adaptability and dynamism demanded by biological processes in a variety of cellular functions, such as gene expression^6^, stress responses^7^, signal transduction^8^, and many others^9–11^.

Prion-like domain proteins, which incorporate lowcomplexity domains (LCDs)—a type of intrinsically disordered region (IDR)—form liquid-like biomolecular condensates readily^12–14^. At the molecular level, proteins within these liquid-like condensates form percolated networks characterized by weak and transient intermolecular interactions^15–17^. However, these biomolecular condensates can also transition into an irreversible solidlike state or form fibrils upon progressive strengthening of intermolecular interactions, which increases the structural rigidity of the assemblies^18–21^. For LCDs containing low-complexity aromatic-rich kinked segments (LARKS), the liquid-to-solid transition may emerge from the formation of stacks of inter-protein cross-*β*-sheets that yield amyloid-like structures^20,22^. Whether a LARKS-containing condensate displays liquid-like or solid-like properties is influenced by several factors, such as temperature^26^, pH^27,28^, salt gradients^29,30^, or even intrinsic changes across the amino acid protein sequence^31^. Specifically, mutations and post-translational modifications are key regulators of the properties of these condensates ^31,32^. For instance, precise mutations in diverse RNA-binding proteins (RBPs) containing LCDs, such as FUS^33,34^, TDP-43^25^, or hnRNPA1^35,36^ can significantly trigger their liquid-to-solid transition. Notably, mutations that favour such transitions have been identified in amyotrophic lateral sclerosis (ALS) and frontemporal dementia (FTD) patients^25,37,38^. Similarly, mutations in *α*-synuclein—a protein associated with Parkinson’s disease^39,40^—and in the Alzheimer-related Tau protein^41^ promote their phase separation into liquid-like condensates, and subsequent ageing into solid-like assemblies through the formation of harmful amyloids.

The dysregulation of liquid-like FUS protein condensates has significant implications for both normal cellular function and disease. On the one hand, FUS plays a key role in stress granules and paraspeckles^6^, and it is also involved in various biological processes, including RNA and DNA metabolism^42^, noncoding RNA regulation^43^, and ribosome biogenesis^44^. On the other hand, dysregulation of FUS condensates is linked to neurodegenerative disorders^34,45,46^, particularly ALS^34,37,47^ and FTD^26,45^. This ambivalence is influenced by specific amino acid mutations. Indeed, certain FUS mutations, such as those in its low-complexity domain (e.g., G156E ALS mutant^46,48^) and the RNA-recognition motif (RRM)^49^, induce abnormal phase behavior. Studies have shown that cation-*π* interactions between tyrosine (Y) residues in the LCD and arginine (R) residues in the RRM and arginine-glycine-rich regions (RGGs) drive the formation and stability of FUS condensates, as well as their aberrant liquid-to-solid transition into insoluble phases^48–50^. Additionally, other amino acid interactions contribute to both the phase separation capacity of FUS and its progressive hardening^51,52^. In disease, FUS forms fibrils and cross-*β*-sheet stacks involving multiple LARKS within its LCD^20^. Specific mutations can also alter hydrogen bonding, hydrophobic interactions, and salt bridges^53–56^, which crucially control its phase behavior. Importantly, glycine (G) residues enhance the fluidity of condensates, while glutamines (Q) and serines (S) promote progressive solidification through steric-zipper transitions^34,57,58^.

A comprehensive study of the intermolecular interactions that regulate the thermodynamic stability and material properties of FUS condensates is challenging to achieve through experimental techniques on its own^59–61^. While techniques such as fluorescence recovery after photobleaching (FRAP), fluorescence spectroscopy, and microrheology (both active and passive)^29,34,62,63^ provide valuable insights into condensate ageing, they are often insufficient to fully uncover the molecular mechanisms underlying these transitions. In this context, computational methods are crucial for interpreting experimental data, achieving sub-molecular resolution, and characterizing interactions and molecular ensembles of the condensate individual constituents^60,64–68^. Previous computational studies have examined the impact on the stability and structure of mutations like R521C, R521H, and P525L in the arginine-glycine-rich (RGG3) region of FUS^49^. Similarly, coarse-grained models have explored the effect of phosphorylation on FUS condensate architecture^69^ and the molecular mechanisms of RNA recognition influenced by mutations in the KK-loop of the RRM region^55^. However, critical information about how specific amino acid mutations across the FUS low-complexity domain—which contains multiple aggregation-prone segments—affect condensate material properties and ageing is still missing.

Residue-resolution coarse-grained models have surfaced as powerful tools that can bridge the physicochemical features of individual proteins to their impact on the phase behaviour at condensate level^70–74^. In this work, we use the Mpipi-Recharged model^75^ to investigate the impact of specific mutations on condensates formed by either the LCD of FUS or the full FUS protein, both of which are prone to develop *β*-sheet accumulation and aberrant solidification over time^17,76–78^.

The Mpipi-Recharged model drastically improves the description of charge effects in biomolecular condensates, while maintaining the excellent predictions for other systems (i.e., prion-like domains^79^) of its predecessor (Mpipi force field^70^)^75,80^. Importantly, Mpipi-Recharged still considers solvation effects implicitly for computational efficiency. We explore the impact of discrete mutations across the FUS-LCD on the rate of condensate ageing; defined as the average incubation time for observing the formation of the first cross-*β*-sheet structure. A number of sets of mutations are tested by substituting polar residues (G, S, P, Q, or T) at various positions along the LCD to either positively charged (K, R), negatively charged (E, D), or non-polar (A) amino acids, while preserving the integrity of key LARKS motifs (i.e., ^37^SYSGYS ^42, 54^SYSSYGQS^61^ and ^77^STGGYG ^82^). This selection allows for a comprehensive assessment of how electrostatic and hydrophobic interactions influence the ageing kinetics of initially liquidlike condensates. Additionally, it provides insights into how changing the identity and position of specific amino acids affects condensate phase behaviour. To model condensate ageing, we incorporate a non-equilibrium algorithm^17^ into our Molecular Dynamics (MD) simulations of the Mpipi-Recharged model^75^. This algorithm enforces inter-protein structural transitions via non-conservative forces informed from binding free energies derived from all-atom simulations^16^. Our approach allows us to investigate the impact of sequence variations on the phase diagram and the time-dependent material properties of biomolecular condensates as they age.

## RESULTS AND DISCUSSION

### A. Phase diagram and time-dependent material properties of FUS and FUS-LCD condensates

To elucidate the impact of sequence mutations on FUS ageing kinetics, we first need to benchmark the phase diagram and viscoelastic properties of the wildtype sequence (see section SII in Supplementary Material, SM). We model both FUS and FUS-LCD using the Mpipi-Recharged force field^75^, a residue-resolution coarse-grained model developed by us. By using a pair-specific asymmetric Yukawa electrostatic potential to describe charge–charge interactions, Mpipi-Recharged improves the description of charge-driven effects in protein phase behaviour. The Mpipi-Recharged model (for technical details on the model potential and parameters please see Section SI in SM) captures subtle charge effects that are difficult to describe without explicit solvation, such as the impact of charge blockiness, stoichiometry changes, and salt concentration variation in condensate formation^75^. Moreover, it further improves the parameterisation of non-electrostatic forces, enhancing its predictions for the experimental hnRNPA1 benchmark set^31^ with respect to previous models^80^. Mpipi-Recharged also provides enhanced predictions of phase diagrams for other prion-like domains and multi-domain proteins such as hnRNPA1, HP1, DDX4, or R12 among others^75^. The parameterisation of the model has been derived from a combination of atomistic simulations of amino acid pairs^70,75^, bioinformatics data^83^, and the hydrophobicity scale of pure amino acids^72–74,84,85^. In this model, IDRs are described as flexible polymers of one-bead per amino acid linked by a harmonic potential, and globular domains are treated as rigid bodies to maintain their secondary structure according to high-resolution models. The solvent and ions in the medium are treated implicitly for computational efficiency.

To validate the model for studying the phase behaviour of FUS, we first perform Direct Coexistence (DC) simulations^86–88^ of both FUS and FUS-LCD to compute their phase diagrams in the temperature–density (*T*-*ρ*) plane (Fig. 1a) (see section SIII for details on this calculation). Consistent with experimental findings^14^, FUS condensates exhibit significantly greater stability compared to the LCD alone. Notably, the critical solution temperature (*T*_*c*_) of FUS is more than 35 K higher than that of its LCD. We compare the saturation concentration (*c*_*sat*_) predicted by the Mpipi-Recharged model for both FUS-LCD and FUS condensates with the experimental *c*_*sat*_ values reported in Refs.^30,82,89^ (Fig. 1a (inset)). Remarkably, we find an excellent agreement between the model predictions and *in vitro* measurements, showing that FUS-LCD requires almost 20 times higher saturation concentration to undergo phase separation^33,51^ compared to the full FUS sequence.

**FIG. 1.**
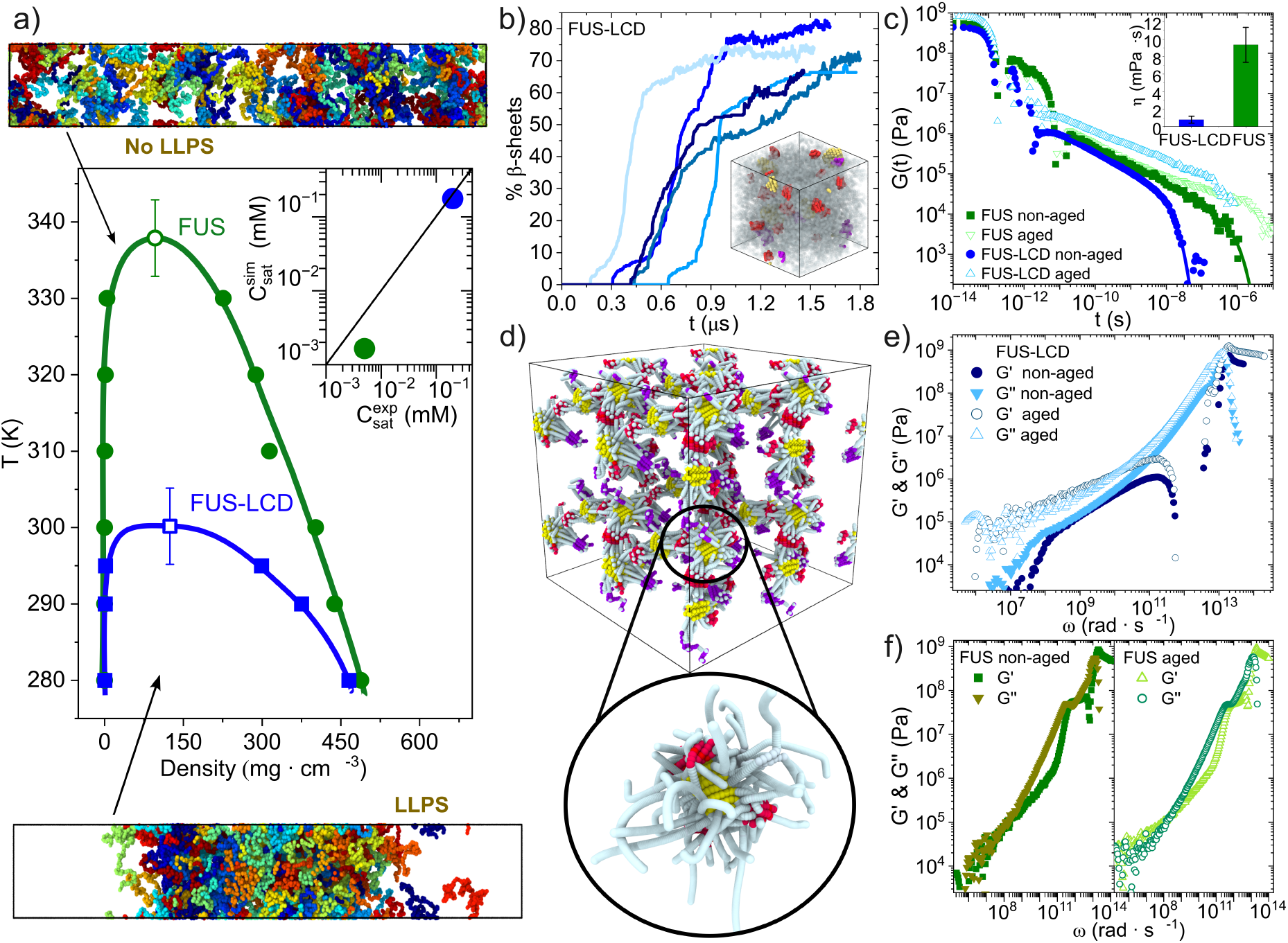
Characterization of FUS and FUS-LCD coexistence lines and time-dependent condensate viscoelastic properties. a) Phase diagram in the *T* –*ρ* plane for FUS (green) and FUS-LCD (blue). Filled symbols represent the coexistence densities obtained via Direct Coexistence simulations, and empty symbols depict the estimated critical points through the laws of critical exponents and rectilinear diameters^81^. Inset: Predicted saturation concentrations from Mpipi-Recharged simulations vs. in vitro experimental values for FUS (green) and FUS-LCD (blue)^30,82^. Representative snapshots of a simulation slab of FUS under LLPS conditions (Bottom) and above the critical solution temperature (Top), where each protein replica is depicted by a different colour. b) Time-evolution of the inter-protein *β*-sheet concentration (expressed in %) for different trajectories of FUS-LCD condensates at 290 K. Inset: Rendered image of the bulk condensate where the inter-protein *β*-sheet structures are highlighted in yellow, red, and purple, according to their corresponding sequence, and the remaining protein residues are coloured in grey. c) Shear stress relaxation modulus *G*(*t*) of FUS (green) and FUS-LCD (blue) at 290 K and condensate bulk conditions. Filled symbols indicate measurements before the emergence of cross-*β*-sheets, and empty symbols after an incubation time of 2 *μ*s. Inset: Viscosity values for non-aged condensates of FUS-LCD and full FUS determined from the integration of *G*(*t*). d) Inter-protein *β*-sheet network connectivity of aged FUS-LCD condensates after 1.5 *μ*s of incubation time as computed through the Primitive Path Analysis. e) Storage *G*′ and loss *G*′′ moduli of FUS-LCD condensates before and after condensate ageing (2 *μ*s) in bulk conditions at 290 K. f) *G*′ and *G*′′ of FUS condensates before and after condensate ageing.

Since structural transitions of LARKS into cross-*β*-sheets can induce progressive solidification of FUS condensates^16,20,90,91^, we now simulate condensate ageing by coupling our dynamic algorithm^16,17,92^ to the Mpipi-Recharged model. This algorithm accounts for changes such as strengthening of inter-protein interactions, local protein rigidification, and shifts in the intermolecular organisation due to the irreversible accumulation of interprotein *β*-sheets in a time-dependent manner, and as a function of the local protein concentration (see Section IV of the SM for details). As previously demonstrated through atomistic potential-of-mean-force calculations^16^, the binding interaction strength during the disorder-to-order transition of the LARKS in the LCD of FUS can increase by up to an order of magnitude compared to the intrinsically disordered state. Such stronger LARKS cross-*β*-sheet interactions within the LCD can drive the condensate into more viscous phases, and eventually into gel-like or even glassy states^29,91^. In Fig. 1b, we show the time-evolution of inter-protein *β*-sheet formation (expressed in % with respect to the maximum number of inter-protein *β*-sheets which can be formed) for five independent trajectories of FUS-LCD condensates in bulk conditions at 290 K. For all condensates, after a certain nucleation lag time (in average ~ 0.4 *μ*s), we observe the formation of inter-protein *β*-sheets. We include a rendered image of an aged condensate in the inset of Fig. 1b, highlighting the cross-*β*-sheet clusters formed in yellow, red and purple depending on the specific LARKS sequence involved in the cross-*β*-sheet assembly, while the rest of the protein residues are shaded in grey.

We examine the material properties of FUS-LCD and FUS condensates in both aged and non-aged states. In the limit of small mechanical perturbations, the viscoelastic behaviour of condensates is characterized by the stress relaxation modulus *G*(*t*)^93^. The shape of the time evolution of the stress relaxation modulus, *G*(*t*), exhibits two distinct regimes: a short-time relaxation region, dominated by fluctuations in short-range interactions and internal conformational rearrangements; and a long-time regime, where the curvature of the function is dictated by the relaxation of intermolecular forces, largescale conformational changes, and the translational diffusion of proteins within the condensate. To compute *G*(*t*),

we perform simulations in the canonical ensemble (e.g., at constant number of particles, volume, and temperature) at 290 K for both FUS and FUS-LCD condensates (Fig. 1c) in bulk conditions (see Section VI of the SM for technical details on these calculations). The results for non-aged condensates (i.e., in absence of inter-protein *β*-sheets; filled symbols) show the characteristic liquid-like behaviour of a polymer melt, defined by a continuous and sharp decay of *G*(*t*) within the simulation timescale. The area under the curve represents the zero-shear rate viscosity, 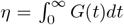, which we show in Fig. 1c (inset). Although the Mpipi-Recharged model cannot provide absolute viscosity values due to its coarse-grained nature and the use of an implicit solvent^94,95^, our results show that the predicted relative viscosities for non-aged FUS and FUS-LCD condensates align with experimental trends. Specifically, the viscosity of FUS condensates is predicted to be an order of magnitude greater than that of the FUS-LCD condensates. In vitro, reported values for FUS condensates range from 3–7 Pa · s^90,96^, while the viscosity for its LCD is approximately 0.4–0.9 Pa · s^33,97^. In particular, the viscosity ratio predicted by the Mpipi-Recharged model, *η*_*FUS*_/*η*_*FUS−LCD*_ ~ 12 (Fig. 1c (inset)), is fully consistent with this experimental difference.

We now investigate the viscoelastic behaviour of aged condensates by monitoring the time-evolution of *G*(*t*) (Fig. 1c). The significantly reduced slope of *G*(*t*) (Fig. 1c; empty triangles) and the absence of a noticeable decay in the function as compared to the pre-aged condensates (filled symbols) indicates that proteins are kinetically trapped, evidencing a liquid-to-solid transition. This is consistent with the fact that the concentration of interprotein *β*-sheets within the condensates is over 70% of the total cross-*β*-sheets which can be formed (Fig. 1b). To elucidate how the topology of the intermolecular network prevents translational diffusion in aged assemblies, we now perform a Primitive Path Analysis (PPA)^17,98–100^ calculation for an aged FUS-LCD condensate simulated over 1.8 *μ*s (see Section SVII in the SM). PPA operates under the following key assumptions: (i) the locations where inter-protein *β*-sheets are bound to each other are fixed in space, (ii) bonds between consecutive residues are treated with a zero equilibrium bond length, and (iii) the intramolecular excluded volume is set to zero. The algorithm then minimizes the contour length of the protein domains connecting structured LARKS while maintaining the topology of the underlying network^100^. In Fig. 1d, the PPA of an aged FUS-LCD condensate is shown. The system has been replicated in the 3-dimensional space for improving the visualization of the underlying network. An elastically percolated network of cross-*β*-sheet clusters (coloured in purple, red, and yellow according to the specific LARKS sequence) emerges frustrating the protein diffusion and driving condensate kinetic arrest.

We also compute the storage (elastic, *G*′) and loss (viscous, *G*′′) moduli for both FUS-LCD (Fig. 1e) and FUS (Fig. 1f) condensates as the real and imaginary components of the Fourier transform of *G*(*t*). In non-aged assemblies, at low frequencies (*ω*), the elastic response, *G*′, remains lower than the viscous response, *G*′′, and the phase angle *δ* = arctan(*G*′′*/G*′) is approximately *π/*4. This behaviour is characteristic of a liquid-like viscous phase, where intra- and inter-molecular interactions have sufficient time to relax and dissipate energy. In contrast, for aged condensates, *G*′ is higher than *G*′′ in the low frequency regime, in consonance with the PPA calculation shown in Fig. 1d. In aged condensates, as frequency decreases, we observe a crossover (*G*′ = *G*′′) marking the transition to a solid or glassy state where molecular rearrangements are too slow to dissipate energy instead of storing it. These results are consistent with the continuous increase in viscosity observed for FUS^29,90^ and FUS-LCD condensates in vitro^91^, which has been attributed to the progressive accumulation of cross-*β*-sheet content as the protein becomes highly concentrated.

### B. Charged residue mutations in FUS-LCD critically control condensate ageing

FUS is a multi-domain protein with a sequence of 526 amino acids and two globular domains; an RNA-recognition motif (RRM) and a zinc finger (ZF; see Fig. 2a). FUS possesses three RGG regions, whose interactions with the LCD are thought to be the main driver for phase separation^52,77,94^. Importantly, several pathogenic glycine mutations across the LCD and adjacent regions (e.g., G191S, G225V, G230C, G156E, G187S, G225V^34,48,58,101,102^) have been shown to disrupt the liquid-like behaviour of FUS condensates by accelerating its liquid-to-solid transition. To further explore these mechanisms, we examine the impact of specific amino acid mutations within FUS-LCD on controlling its propensity to develop inter-protein *β*-sheet transitions (Fig. 2a). We designed a set of mutant sequences in which none of the residues within the three LARKS motifs in the LCD (i.e., ^37^SYSGYS ^42, 54^SYSSYGQS^61^ and ^77^STGGYG ^82^) were altered. Instead, mutations were introduced at positions adjacent to these LARKS (highlighted in red, Fig. 2a). Hereafter, we adopt the notation Mutant *n*-*X*, where *n* indicates the mutant type (1–4, Fig. 2a) and *X* represents the amino acid substituted relative to the wild-type sequence.

**FIG. 2.**
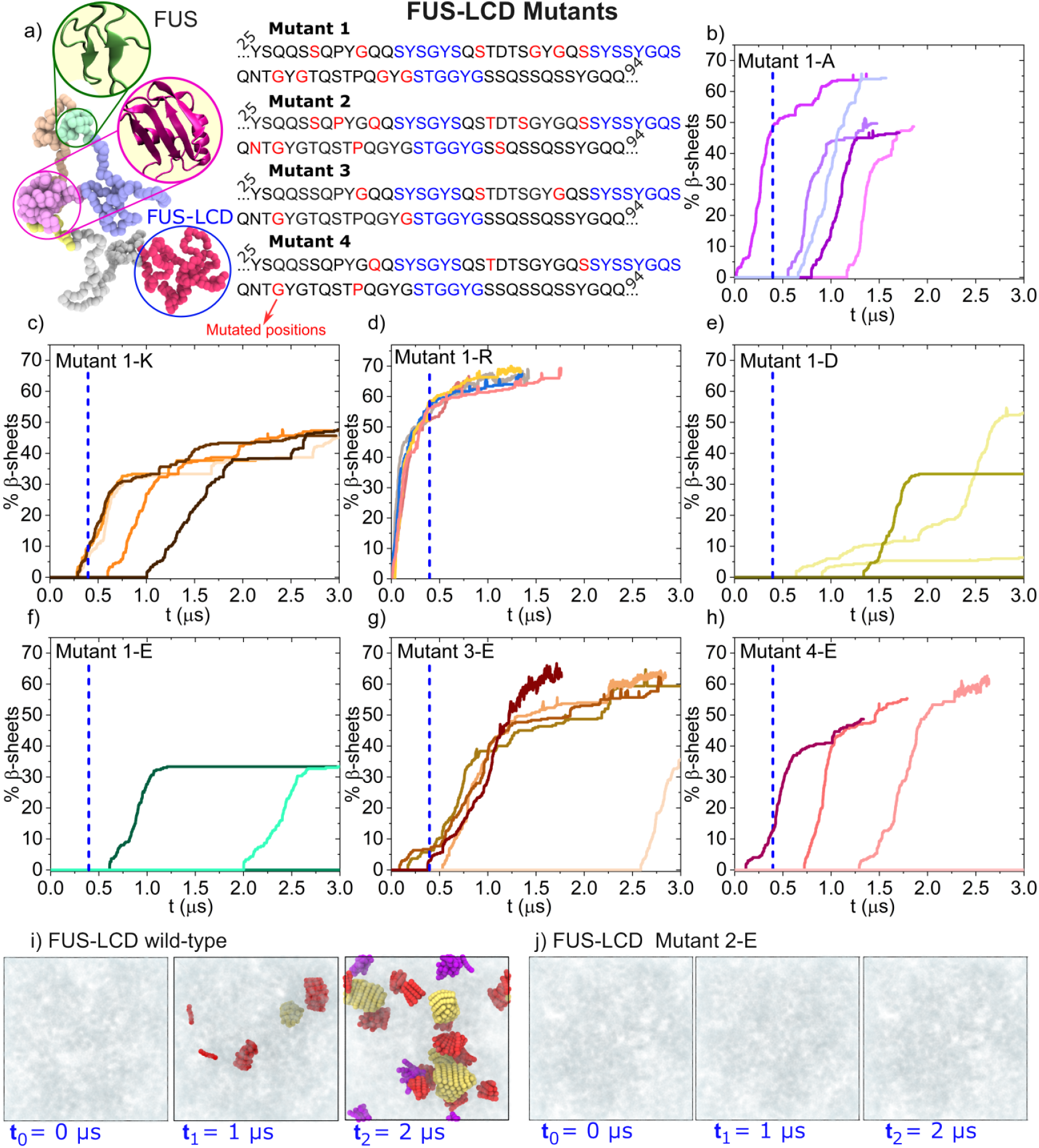
Mutations in FUS-LCD heavily regulate the onset of inter-protein *β*-sheet transitions in condensed phases. (a) Scheme of the FUS sequence colouring the different domains as LCD (red), RGG1 (gray), RRM (pink), RGG2 (light orange), ZF (green), and RGG3 (violet). The four types of mutant substitutions on the FUS-LCD sequence are provided on the right, marking in red the specific residues which aim to be mutated, and in blue the protein LARKS. (b-f) Time-evolution of inter-protein *β*-sheet transitions (in percentage) for 5 independent condensate trajectories of Mutant 1 with different substitutions to: alanine (b), lysine (c), arginine (d), aspartic acid (e), and glutamic acid (f). The same results but for Mutant 3 and 4 using glutamic acid substitutions are shown in panels (g) and (h), respectively. A vertical dashed blue line indicates the average nucleation time for the wild-type FUS-LCD sequence, *τ*_*wt*_. Representative snapshots of the timeevolution of FUS-LCD (i) and FUS-LCD Mutant 2-E (h) condensates highlighting the cross-*β*-sheet structures formed by the three LARKS (red, yellow, and purple).

By designing targeted sets of mutations, we can systematically explore how mutation position, number of mutated residues, and identity influence the kinetics of inter-protein *β*-sheet transitions via specific physicochemical interactions. We classify our mutants as follows: **Mutant 1**, where 10 glycine and serine residues at positions marked in red in Fig. 2a are mutated; **Mutant 2**, with mutations at 10 different positions—including residues glycine, serine, proline, glutamine, threonine, and asparagine—to probe the impact of both chemical identity and sequence location compared to Mutant 1; and **Mutants 3 and 4**, containing half the mutations described in Mutant 1 and Mutant 2, respectively. We first investigate how the chemical nature of substituted amino acids affects FUS-LCD ageing kinetics by introducing charged residues—positively charged (arginine and lysine) and negatively charged (glutamic acid and aspartic acid)—anticipating that electrostatic repulsion may disrupt local LARKS high-density fluctuations. Additionally, we test mutations to alanine, a hydrophobic residue, as a neutral control. The original amino acids replaced in these mutants (glycine and serine) are neutral hydrophobic residues commonly found in phase-separating RNAbiding proteins^94^, such as FUS (see Fig. 2a).

For all mutants, we simulated the dynamic formation of cross-*β*-sheet assemblies using five independent trajectories (with different initial velocities) of FUS-LCD condensates, each containing 100 proteins. Simulations were performed at the equilibrium density of FUS-LCD condensates at *T* = 290 K (i.e., *T/T*_*c*_ ~ 0.97, Fig. 2b-h; see Section SIV for detailed methods). Notably, mutations in Mutant 1 significantly influence FUS-LCD ageing kinetics. To compare the onset of cross-*β*-sheet formation directly between the wild-type and mutants, we include the average nucleation time (*τ*_*wt*_) of wild-type FUS-LCD (dashed blue line in Fig. 2b-h), defined as the average time at which the first cross-*β*-sheet appears.

Mutant 1-A exhibits moderately delayed nucleation relative to the wild-type FUS-LCD (Fig. 2b), with most trajectories initiating cross-*β*-sheet formation later than *τ*_*wt*_ (Fig. 1b). Because alanine is slightly larger than glycine^70,75^, and both act as a spacer (i.e., increasing protein solubility and forming weak residue–residue interactions)^31^, mutations from glycine to alanine hinder cross-*β*-sheet formation. Nonetheless, both wild-type FUS-LCD and Mutant 1-A condensates reach concentrations exceeding 50% LARKS participation in cross-*β*-sheets. Recent simulations indicate that cross-*β*-sheet fractions above 20% significantly alter condensate material properties, inhibiting condensate fusion and promoting multiphasic architectures even in single-component protein condensates^103^.

We next substituted neutral residues with positively charged residues (e.g., arginine and lysine; Fig. 2c-d) at the positions defined by Mutant 1. Mutant 1-K (with increased lysine content) condensates display slower nucleation times than the wild-type, whereas Mutant 1- R (with increased arginine content) transitions prematurely into ordered *β*-sheets. This notable difference arises from strong cation-*π* interactions between arginine and the multiple aromatic residues in the LCD, which increase condensate stability and density^52,104^, and in turn, enhance ageing kinetics. Additionally, recent atomistic simulations have shown that lysines experience stronger electrostatic self-repulsion than arginines^75^, resulting in more effective disruption of high-density LARKS fluctuations in Mutant 1-K compared to Mutant 1-R. Arginine acts as a context-dependent sticker, forming strong interactions with aromatic, polar, and negatively charged residues^14,31,32,50,79^, whereas lysine interactions are weaker and shorter-lived^30,75,104^. Consequently, although both residues are positively charged, their influence on FUS-LCD ageing kinetics significantly differs. These findings align with previous coarse-grained simulations using the HPS-Cation-*π* model^73^, where arginine substitutions in FUS-LCD were similarly found to accelerate inter-protein *β*-sheet formation^50^. Remarkably, Mutant 1-R not only accelerates nucleation but also increases the growth rate and total number of cross-*β*-sheets formed compared to Mutant 1-K.

When neutral amino acids are substituted by negatively charged residues (Mutants 1-E and 1-D; Figs. 2e-f), the formation of cross-*β*-sheet transitions is significantly hindered. Unlike positively charged mutations, substitution with aspartic or glutamic acid in the LCD do not readily promote intermolecular contacts with neighbouring aromatic residues, as observed in Mutant 1-R. Moreover, electrostatic self-repulsion between the negatively charged side chains of aspartic acid or glutamic acid further disrupts local high-density concentrations of LARKS, thereby frustrating their transition into cross-*β*sheets (Figs. 2e-f). Although both aspartic and glutamic acid delay the onset of ageing, glutamic acid shows a slightly greater efficacy in preventing structural transitions. In Mutant 1-D condensates, two of the five simulation trajectories show no inter-protein *β*-sheet formation. In contrast, for Mutant 1-E condensates, three out of five trajectories remain devoid of structural transitions for over 3 *μ*s. Furthermore, when cross-*β*-sheets do emerge in Mutant 1-E, their formation occurs on longer timescales compared to Mutant 1-D.

Interestingly, the growth of cross-*β*-sheets in both Mutant 1-E and Mutant 1-D is frustrated at significantly lower concentrations (i.e., below 50%) compared to all the other studied mutants in this work, including the wild-type sequence. This suggests that aspartic acid and glutamic acid substitutions not only preclude inter-protein *β*-sheet formation but also mitigate ageing-related effects, such as kinetic arrest, in the material properties of the condensates. Our simulations indicate that the superior effectiveness of glutamic acid mutations preventing cross-*β*-sheet transitions compared to aspartic acid is possibly due to its larger excluded volume, stronger self-repulsion^75^, and the weaker associations it forms with positively charged species, which more effectively suppress high-density LARKS fluctuations when substitutions occur near LARKS regions.

To further investigate the role of residue abundance and sequence patterning in the glutamic acid variant (Mutant 1-E), we simulated condensates of Mutants 2-E, 3-E, and 4-E (Fig. 2g–j). The rate of disorder-to-order transitions in Mutants 3-E and 4-E is markedly higher than in Mutants 1-E and 2-E, respectively, owing to the lower number of glutamic acid substitutions (5 instead of 10). Remarkably, none of the simulation trajectories for Mutant 2-E exhibit cross-*β*-sheet transitions over the full 3*μ*s duration, underscoring the critical importance of specific substitution positions in modulating fibrillization (Fig. 2i–j). In Mutant 3-E, the five substituted residues consist of one serine and four glycines, and the average nucleation time for cross-*β*-sheet formation (Fig. 2g) is comparable to that of FUS-LCD (wild-type) condensates. In contrast, the ageing kinetics of Mutant 4-E is substantially slower than that of Mutant 3-E, primarily due to the different identities and positions of the substituted residues—which include serine, glycine, glutamine, threonine, and proline (Fig. 2a). Notably, even with only five substitutions, 40% of the Mutant 4-E condensates show no structural transitions over the course of the simulation. These results highlight that, beyond the number of mutations, the identity and sequence location of the substituted residues play a pivotal role in controlling the liquid-to-solid transitions of condensates.

Figures 2i and 2j display representative snapshots from a FUS-LCD and a Mutant 2-E condensate trajectory over time, with cross-*β*-sheet assemblies shown in yellow, red, and purple. Strikingly, Mutant 2-E condensates fail to form cross-*β*-sheets within the same simulation timescale in which FUS-LCD accumulates high concentrations of these assemblies (Fig. 1b). We attribute the stronger inhibition of ageing kinetics in Mutant 2-E relative to Mutant 1-E to the broader diversity of substituted residues. While Mutant 1-E includes only glycine and serine substitutions, Mutant 2-E incorporates substitutions of serine, glycine, proline, glutamine, asparagine, and threonine.

The Mpipi model used in this work, which has demonstrated near-quantitative accuracy in predicting experimentally measured condensate stability and material properties^31,75,^, enables us to probe the molecular mechanisms by which specific mutations alter condensate behaviour. According to our simulations, the substituted residues in Mutant 2-E are capable of engaging in hydrophobic interactions with other proteins in the condensate and form preferential contacts with glycine or serine. The introduction of glutamic acid in these mutants replaces associative interactions that typically favour LARKS–LARKS contacts with electrostatic repulsion. This substitution disrupts the formation of high-density local concentrations of LARKS, thereby inhibiting cross-*β*-sheet formation. Overall, our findings highlight the critical influence that even single-residue mutations can exert on protein liquid-to-solid transitions. This is fully consistent with the experimentally observed G156E mutation, which reliably increases both the viscosity and ageing kinetics of FUS condensates, promoting their maturation into solid-like states^33^.

We now compare the kinetics of inter-protein *β*-sheet formation across all FUS-LCD mutants in Fig. 3a-b. To this end, we calculate the average nucleation time (*< τ >*) for the formation of the first cross-*β*-sheet nucleus by averaging over all independent trajectories for each condensate variant shown in Fig. 2. Since protein aggregation and condensate ageing kinetics driven by cross-*β*-sheet transitions can be approximated as a first-order reaction^17,50^, we estimate *< τ >* for the FUS-LCD mutants in which some of the trajectories did not exhibit any disorder-to-order transition using a firstorder kinetic expression (see Section SV in the SM for details).

**FIG. 3.**
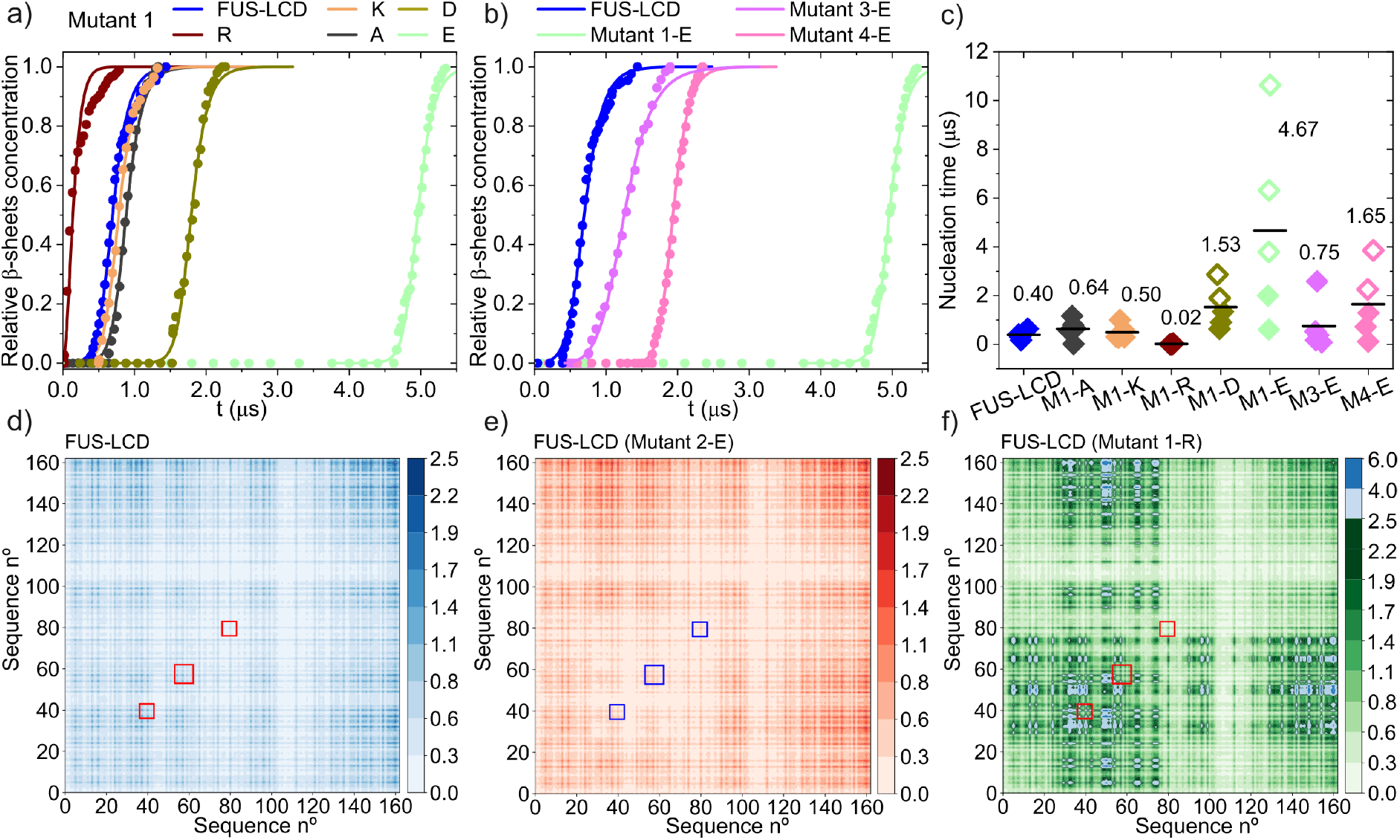
Ageing kinetics and intermolecular contact maps of FUS-LCD mutants. (a) Normalised number of interprotein *β*-sheet transitions as a function of time averaged over five different independent trajectories for FUS-LCD and Mutant-1 variants with alanine (A), arginine (R), aspartic acid (D), lysine (K), and glutamic acid (E). Solid lines are included as a guide to the eye. (b) Normalised number of inter-protein *β*-sheet transitions as a function of time averaged for different FUS-LCD variants only including glutamic acid mutations. (c) Average nucleation time (*< τ >*, black horizontal bar) of the different variants to develop inter-protein *β*-sheet transitions. Solid diamonds represent the nucleation time from individual trajectories directly observed from our simulations while empty diamonds are estimates assuming a first-order kinetic reaction. (d–f) Intermolecular contact map probability (expressed in residue-residue % contact frequency) prior ageing for FUS-LCD (wildtype) (d), Mutant 2-E (e), and Mutant 1-R (f) condensates. The squares included in each contact map indicate the three different LARKS identified across the FUS-LCD sequence^20^.

The advantage of this approach is that it circumvents the need to wait for all trajectories to undergo structural transitions—an important consideration, as several mutants display transition times beyond the accessible simulation window (i.e., *>* 4 *μ*s; Fig. 3c). In Fig. 3c, we report the average nucleation time (⟨*τ*⟩) of cross-*β*-sheet assemblies for all FUS-LCD variants. Filled diamonds indicate nucleation times directly measured within the simulation timescale (i.e., *<* 3 *μ*s), while empty diamonds represent estimates obtained through the first-order kinetics model (see the benchmarking and validation approach in Section SV of the SM).

Our simulations reveal that substitutions with negatively charged residues, such as the Mutant 1-E, delay the nucleation time of cross-*β*-sheet clusters by approximately an order of magnitude compared to the wild-type FUS-LCD sequence. For Mutant 2-E, the extent of this kinetic deceleration could not be quantified, as none of the five simulated trajectories exhibited interprotein structural transitions within the accessible simulation time scales. Remarkably, glutamic acid substitutions appear more effective than aspartic acid substitutions in frustrating disorder-to-order transitions even for the same substituted positions across the sequence (e.g., Mutant 1-E vs. Mutant 1-D). Similarly, aspartic acid substitutions are more effective than either lysine or alanine at disrupting ageing, while arginine substitutions promote, rather than hinder, ageing kinetics.

To further analyse the impact of specific mutations on FUS-LCD condensate ageing kinetics, we represent the normalised average time-evolution of the cross-*β*-sheet concentration^106^ for the different variants (Fig. 3a-b). We average *< τ >* from the different independent trajectories. The cross-*β*-sheet growth is obtained by averaging the interval from the nucleation time onwards in the trajectories that displayed cross-*β*-sheet transitions over time (see Section SV of SM). In Fig. 3a, we plot the resulting curves for all the variants corresponding to Mutant 1. This figure highlights how inter-protein *β*sheet transitions heavily depend on the identity of the substituted residues. While Mutant 1-R condensates (brown curve; Fig. 3a), display the fastest accumulation of cross-*β*-sheet fibrils—due to the strong associative interactions that arginine forms with aromatic and negative residues^31,50,107,108^—lysine and alanine mutants approximately yield the same ageing kinetics of the wildtype sequence (blue curve). Importantly, despite lysine being able to establish intermolecular interactions of the same type as arginine (i.e., electrostatic associations and cation-*π* interactions^30^), the formation of inter-protein structural transitions for Mutant 1-R and 1-K is radically different due to the lower affinity of lysine for aromatic and polar residues^31,32,107,109^ as well as the higher electrostatic repulsion between lysine–lysine compared to arginine–arginine pairs ^63,75,110^.

Our simulations suggest that introducing negatively charged residues—as in Mutants 1-D and 1-E—is the most effective strategy for disrupting high-density local fluctuations of LARKS that drive inter-protein *β*-sheet formation. Strikingly, mutants incorporating glutamic acid substitutions (light green curve in Fig. 3a) delay ageing kinetics by approximately an order of magnitude compared to the wild-type FUS-LCD sequence. This is consistent with recent experimental findings showing that the inclusion of aspartic or glutamic acid residues into protein sequences hinders phase separation^31,32^, largely due to their disruptive effect on stabilising intermolecular contacts^31^. However, the growth dynamics of cross*β*-sheet clusters following the formation of the initial nucleus appear to be only marginally affected by mutations (Fig. 3a). This apparent insensitivity arises from the normalisation of cross-*β*-sheet concentration. As shown in Fig. 2b–h, the actual percentage of LARKS converting into cross-*β*-sheets varies substantially depending on the specific mutations introduced.

A comparison of the average time evolution of cross-*β*-sheet content across all glutamic acid mutant sequences (Fig. 3c) reveals the crucial role of both the number and the positions of the substitutions. For example, the difference between Mutant 1-E and Mutant 3-E highlights how a reduced number of mutations leads to significantly faster structural transitions. Additionally, the identity of the original residues being replaced strongly influences the condensate’s liquid-to-solid transition propensity. While none of the Mutant 2-E condensates exhibit cross-*β*-sheet formation within the accessible simulation timescales, some trajectories of Mutant 1-E do lead to *β*-sheet cluster formation. Similarly, the average nucleation time ⟨*τ*⟩ for Mutant 4-E is twice that of Mutant 3-E, despite both having the same number of mutations in comparable sequence positions. Furthermore, the fact that Mutants 1-E and 3-E, as well as Mutants 2-E and 4-E, share overlapping substitutions allows us to systematically assess the relative impact of specific sequence positions and residue identities on regulating cross-*β*-sheet transitions.

Since mutations play a critical role in modulating the intermolecular interactions that govern the phase behaviour of protein solutions^31,32,80,109^, we next quantify how different substitutions affect the liquid-like network connectivity within FUS-LCD condensates. We focus on the intermolecular contact maps of the wild-type FUS-LCD sequence (Fig. 3d) and the two mutants which display the most contrasting ageing kinetics: Mutant 1-R (Fig. 3f) and Mutant 2-E (Fig. 3e). Intermolecular contact maps for the remaining sequence variants are provided in Section SIX of the SM. Intermolecular contact maps were evaluated prior to the formation of cross-*β*- sheets and under identical temperature and density conditions for all sequences (see Section SVIII of the SM). Analysis of these contact maps within FUS-LCD condensates reveals that the first identified LARKS region (hereafter referred to as LARKS1) exhibits the highest probability of forming intermolecular interactions with the same LARKS region across different protein chains (highlighted by red squares in Fig. 3d), compared to the other two LARKS. The substitutions in Mutant 2-E significantly change the intermolecular interaction maps; specifically, the contact probability of the segment containing the three LARKS (i.e. from the 25th to the 94th residue) decreases by ~ 33% compared to the wild-type FUS-LCD condensate. At the same time, the substitutions enhance the affinity between the LARKS region and the C-terminal domain of the LCD (residues 130–163), which is enriched in glutamine and tyrosine residues. Importantly, this region does not undergo disorder-to-order transitions independently, which is consistent with the inhibition of aggregation. A similar inhibitory effect on LARKS–LARKS interactions is found for Mutants 1-E, 3-E, and 4-E (see Figs. S6-S8 in the SM), which also display a reduced contact probability between LARKS of the same sequence after mutation. In contrast, Mutant 1-R presents a large increase in the LARKS– LARKS contact probability, which is driven by cation- *π* interactions^107,109^ between the mutated arginines and aromatic residues (as shown in Fig. 3f). In Table S3, we quantify the contact probability variation between the three LARKS for the different studied mutants relative to wild-type FUS-LCD. Strikingly, we observe that LARKS–LARKS intermolecular contacts in Mutant 2- E are reduced by up to ~ 50% compared to the wild-type FUS-LCD. Similarly, mutants 1-E, 3-E, and 4-E also show a decrease in the contact probability of LARKS1– LARKS1 of ~ 34%, ~ 21%, and ~ 32%, respectively, which is fully consistent with their relative average nucleation times to develop cross-*β*-sheet fibrils upon incubation (Fig. 3c). On the other hand, Mutant 1-R shows a huge increase of ~ 120% for the same contacts compared to FUS-LCD. These calculations highlight the major impact that charge-residue mutations induce on the intermolecular liquid network connectivity of FUS-LCD condensates, which in turn deeply controls the kinetics of cross-*β*-sheet formation.

### C. Ageing modulation through LCD mutations is transferable to full FUS

The low-complexity domain (LCD) of FUS provides a powerful model system for investigating how specific mutations influence condensate stability and ageing kinetics^34,103,111^. However, in cellular environments, FUS exists as a full-length, multi-domain protein. Thus, to assess whether the effects observed in the isolated LCD translate to the full FUS sequence, it is essential to evaluate how these mutations impact condensate material properties in the context of the entire protein. In this section, we examine how the LCD mutations that most effectively suppressed cross-*β*-sheet formation in FUS-LCD condensates (i.e., Mutants 1-E and 2-E), as well as the mutation that accelerated ageing kinetics (Mutant 1-R), influence the phase behaviour of full-length FUS. In Fig. 4a, we present the phase diagrams (temperature– density plane) of full-length wild-type FUS and the selected mutants. Remarkably, the critical temperatures for all four sequences fall within a narrow range of 3 K: *T*_*c*_ = 340 ± 5 K for Mutants 1-R and 2-E, and *T*_*c*_ = 337 ± 5 K for the wild type and Mutant 1-E. Moreover, the coexistence densities and saturation concentrations required for phase separation are largely comparable across the different sequences as a function of temperature. Only minor differences are observed, with wildtype FUS condensates exhibiting slightly higher equilibrium densities—less than 8% greater—relative to the mutant variants.

**FIG. 4.**
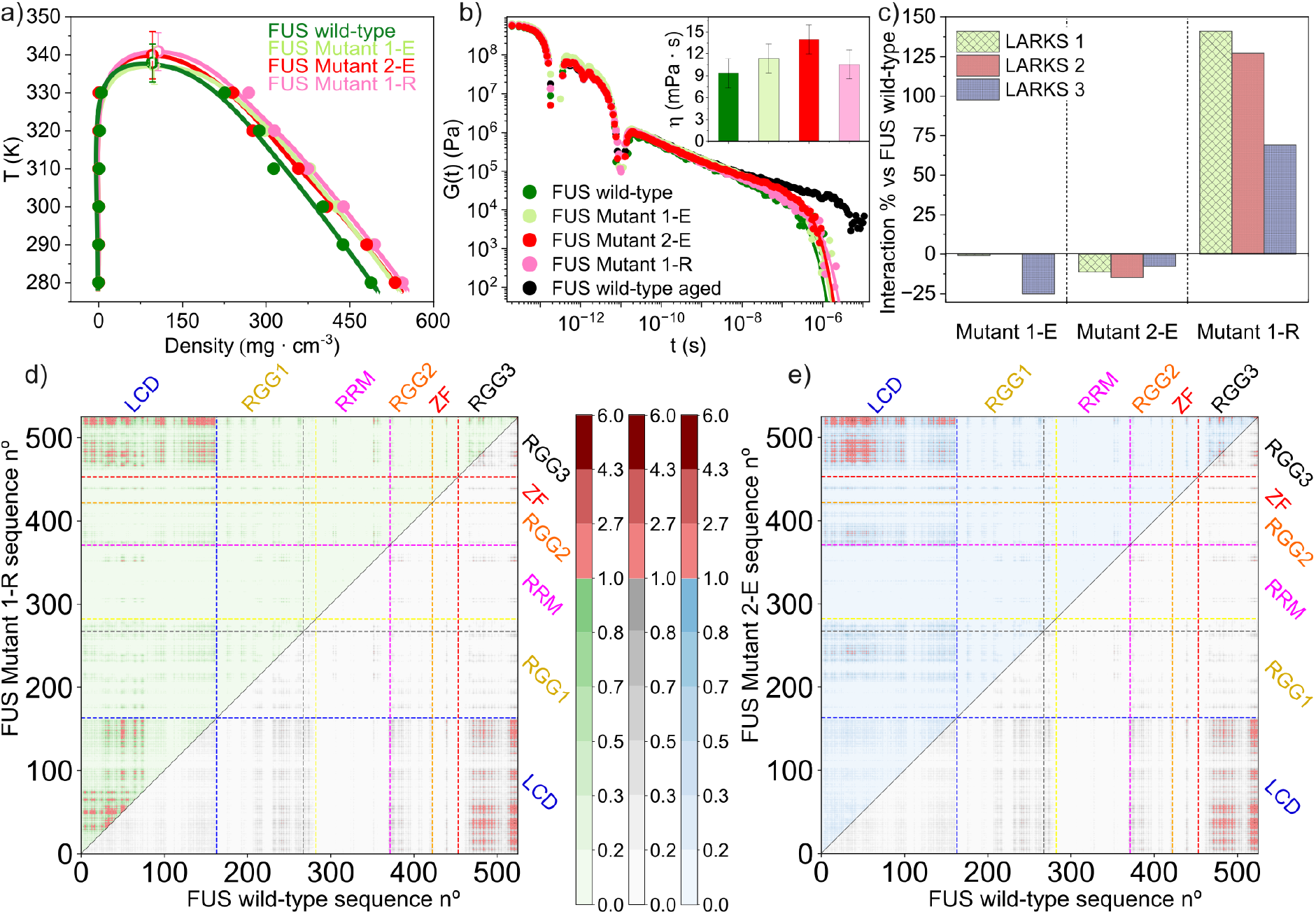
Phase diagram, condensate viscosity, and intermolecular contact frequencies of FUS and E/R mutated variants. a) Phase diagram in the *T* –*ρ* plane for FUS wild-type and Mutants 1-E, 2-E, and 1-R. Filled symbols represent coexistence densities obtained via DC simulations, while empty symbols indicate the estimated critical points. Solid lines are included as a guide to the eye. b) Shear stress relaxation modulus *G*(*t*) at 290 K for non-aged and aged wild-type FUS condensates, as well as for non-aged condensates of Mutants 1-E, 2-E, and 1-R. Solid lines represent Maxwell mode fits to the simulation data. c) Variation of intermolecular LARKS-LARKS interactions (expressed in %) for condensates of Mutants 1-E, 2-E, and 1-R compared to FUS wild-type. d) Map of intermolecular contact frequencies (per protein domain) for condensates of: Mutant 1-R (upper diagonal) vs. FUS wild-type (lower diagonal). e) Map of intermolecular contact frequencies for condensates of: Mutant 1-E (upper diagonal) vs. FUS wild-type (lower diagonal). The scale bar is expressed in percentage of probability of pairwise residue-residue interactions.

We now investigate the impact of the selected mutations on the ageing propensity of the full-length protein. To this end, we perform simulations analogous to those in Fig. 2, at *T* = 290 K and using the respective condensate equilibrium density for each variant (Fig. 4a). Across multiple independent trajectories spanning over 4 *μ*s, we observe no evidence of LARKS disorder-to-order structural transitions into cross-*β*-sheets. This absence of cross-*β*-sheet formation is attributed to the significantly longer timescales required to observe inter-protein *β*-sheet formation in full-length FUS (526 residues) compared to the shorter FUS-LCD segment (163 residues)^90,91,103^. The probability of local high-density fluctuations involving multiple LARKS within a short distance range is markedly reduced when the overall concentration of LARKS in the condensate decreases by approximately a factor of 3.5—for example, three LARKS in 163 residues for FUS-LCD versus three in 526 residues for full FUS.

To qualitatively assess whether the ageing trends observed in FUS-LCD are preserved in full-length FUS, we reparameterise the dynamic algorithm governing LARKS structural transitions^16,17^ (see Section SIV in the SM for details). Specifically, we reduce the coordination threshold required to trigger inter-protein transitions: transitions now occur when four (rather than five) LARKS are detected within the cut-off distance. This adjustment artificially accelerates cross-*β*-sheet formation by bringing the timescale of structural conversion closer to that of protein translational diffusion—despite experimental evidence indicating that structural transitions are typically much slower^90,112^, as reflected in our FUS-LCD results in Figs. 2 and 3. Remarkably, even with this artificially enhanced transition probability, the ageing trends observed for FUS-LCD remain qualitatively consistent in the fulllength protein (Fig. S1). Mutant 1-R exhibits accelerated ageing, with near-instantaneous formation of cross*β*-sheet structures, while Mutants 1-E and 2-E show a markedly slower onset of inter-protein cross-*β*-sheet assembly, mirroring their behaviour in the LCD context.

Since the dynamic formation and dissolution of FUS condensates is essential for their biological function^8,45^, their precise material properties are of critical importance^29^. We, therefore, evaluate the viscoelastic behaviour of FUS and its mutants under pre-ageing conditions—i.e., in the absence of cross-*β*-sheet assemblies and with the ageing dynamic algorithm disabled. Additionally, we assess the viscoelastic response of wildtype FUS after an incubation period of approximately 3.5 *μ*s, using the modified dynamic algorithm parameters described earlier (see Table S2). In Fig. 4b, we present the shear stress relaxation moduli *G*(*t*) for wildtype FUS (both pre- and post-ageing), along with those for Mutants 1-R, 1-E, and 2-E. The corresponding viscosities (inset of Fig. 4b) for all sequences prior to ageing are comparable, ranging from 9 to 12 mPa · s within the uncertainty. These results indicate that, although the introduced mutations significantly modulate the nucleation onset of cross-*β*-sheet transitions, they do not substantially affect the viscoelastic properties of the condensates under pre-ageing conditions. By contrast, wild-type FUS condensates following prolonged incubation exhibit solidlike behaviour (black circles), characterised by a shear stress relaxation modulus that remains nearly constant throughout the simulation timescale, reflecting the formation and persistence of cross-*β*-sheet networks.

To further understand the different ageing propensities that the three FUS mutants display, we now compute the variation (in %) of LARKS–LARKS intermolecular contacts compared to wild-type FUS condensates (Fig. 4c). Similarly to FUS-LCD mutated sequences (Fig. 3d-f), we find that Mutant 1-R displays much higher intermolecular contact probabilities among their LARKS than FUS (over 100% probability for LARKS1 and LARKS2 interprotein contacts). Conversely, Mutant 2-E shows an overall 10% lower probability of LARKS–LARKS interactions for the three sequences, which explains its much lower propensity to develop cross-*β*-sheet transitions over time. Likewise, Mutant 1-E displays a reduced probability for LARKS3–LARKS3 interactions of 25% compared to FUS, although a similar contact likelihood for the two remaining LARKS compared to wild-type FUS (Fig. 4c). Therefore, our contact analysis of LARKS intermolecular interactions for FUS pre-aged condensates supports our findings using only the low-complexity domain of FUS which evidence a significantly lower probability of LARKS high-density local fluctuations upon E mutagenesis.

Since the overall intermolecular liquid network connectivity of the FUS mutants seems to be conserved upon glutamic acid and arginine substitution given their highly similar condensate stability (Fig. 4a) and viscoelastic properties (Fig. 4b), we next compare the full-sequence contact map probability (Fig. 4d-e) of Mutant 1-R (green upper diagonal) and Mutant 2-E (blue upper diagonal) against FUS wild-type (gray scale in lower diagonal). We find that the strong increase of LARKS– LARKS interactions in Mutant 1-R reported in Fig. 4c is mediated by a prominent enhancement of LCD–LCD interactions (amino acids 1-163), which in turn causes a significant reduction in LCD–RGG condensate contacts (Fig. 4d). Such dominance of LCD–LCD interactions with respect to LCD-RGG1, LCD–RGG2, and LCD–RGG3 contacts, which significantly decrease by 20%, 37%, and 12%, respectively, compared to FUS wild-type (see Table S5) facilitate LARKS local high-density concentrations, which rapidly accumulate inter-protein cross-*β*-sheets upon condensate incubation (Fig. S1). In contrast, the contact map probability of Mutant 2-E reveals a reduction in LCD–LCD intermolecular contacts (*>*5%) compared to FUS, and a more prominently increase in LCD–RGG2 contacts (13%) followed by LCD– RGG1 (6%) and LCD–RGG3 (1%) cross-interactions (see Table S5). Negatively charged glutamic acid substitutions within the LCD enhance electrostatic interactions with positively charged residues in the RGG regions. This shift in interaction preference hinders cross*β*-sheet transitions by promoting inter-protein conformational alignments that favour LCD–RGG contacts over LCD–LCD interactions. Overall, our intermolecular contact maps from full-length FUS condensates confirm that the trends observed in FUS-LCD variants regarding their liquid-to-solid transition propensity are preserved in the context of the complete protein sequence.

## I. CONCLUSIONS

In this work, we investigate how specific mutations that increase the number of charged residues within the LCD of FUS regulate the material properties and ageing kinetics of phase-separated condensates. FUS is a key RNA-binding protein involved in multiple cellular processes, including RNA and DNA metabolism, and the dysregulation of its condensates into solid-like assemblies has been linked to neurodegenerative disorders such as ALS and FTD^26,45^. To explore the molecular determinants of this behaviour, we perform residue-resolution simulations using the Mpipi-Recharged model^75^, which has demonstrated excellent performance in recapitulating the effects of single-point mutations on experimental protein phase diagrams^80^. We couple Mpipi-Recharged with a non-equilibrium dynamic algorithm that we developed to model the formation of inter-protein *β*-sheets in protein condensates^16,17^. Our simulations emphasize the excellent agreement of Mpipi-Recharged predictions with experiments, as we show it predicts the saturation concentration of both FUS and FUS-LCD to undergo phase separation (Fig. 1a) and their relative condensate viscosity (Fig. 1b). Furthermore, our simulations also capture the experimentally observed liquid-to-solid transitions in both FUS-LCD^34,91^ and full-length FUS^90,103^ condensates driven by the gradual accumulation of inter-protein *β*-sheets. Crucially, our model enables molecular-level characterisation of the percolated *β*-sheet network that underpins the progressive alteration of condensate viscoelastic properties. The observed cross-over in our simulations between the elastic and viscous moduli—where *G*′ *> G*′′ (Figs. 1e-f)—marks the transition from a liquidlike state to a kinetically trapped, solid-like phase.

To investigate how specific mutations influence condensate ageing, we design a series of substitutions across the low-complexity domain of FUS. These mutations are introduced in residues adjacent to the three experimentally identified LARKS motifs in FUS^20^, while preserving the key amino acids and positions required for LARKS-mediated disorder-to-order transitions. Our results show that substitutions to alanine or lysine have minimal effect on the ageing kinetics of FUS-LCD condensates (Fig. 2b–c). In contrast, mutations to arginine, glutamic acid, or aspartic acid significantly alter the propensity of FUS-LCD to develop pathological cross-*β*-sheet structures. Arginine substitutions strongly promote the formation of ordered *β*-sheets, driven by cation–*π* interactions between arginine residues and aromatic side chains within the LCD^107,109^. These interactions enhance local intermolecular contacts near LARKS-adjacent regions, increasing the likelihood of high-density fluctuations that favour *β*-sheet formation. Conversely, substitutions with negatively charged residues—particularly glutamic acid—effectively suppress cross-*β*-sheet formation (Fig. 2f–j). By analysing the patterning, abundance, and identity of the substituted residues, we find that replacing polar, uncharged side chains such as glutamine, threonine, or serine by self-repulsive residues like glutamic acid leads to a pronounced slowdown in ageing kinetics. Collectively, our simulations highlight the critical role of specific single-point mutations in modulating the onset of condensate hardening and the transition toward solid-like states.

We extend our characterisation of the mutations that impact FUS-LCD condensate ageing most strongly to the full-length FUS protein. Remarkably, we find that the glutamic acid and arginine substitutions introduced in the LCD have minimal impact on the overall phase diagram of FUS, or on the condensate viscosity prior to the accumulation of cross-*β*-sheet structures (Fig. 4a–b). However, the nucleation time preceding the emergence of inter-protein *β*-sheets varies significantly depending on the specific mutations. Faster fibrillation kinetics are associated with a higher frequency of intermolecular LARKS–LARKS contacts formed within each variant’s condensates (Fig. 4c). Notably, the increased LCD–LCD contact probability in Mutant 1-R, which promotes the rapid formation of inter-protein *β*-sheet clusters, also occurs at a much higher rate in the full-length mutant than in wild-type FUS condensates (Fig. 4d). In contrast, negatively charged glutamic acid substitutions, which reduce LCD–LCD interactions and cross-*β*-sheet formation, also decelerate ageing of the full length protein.

Taken together, our non-equilibrium, residue-resolution simulations of biomolecular condensates offer a powerful framework for uncovering how different sets of amino acid mutations regulate the stability and ageing kinetics of a broad range of prion-like protein condensates. Experimentally screening large numbers of sequence variants remains a major challenge^26^; however, our work introduces a computational pipeline through which the effects of sequence mutations on condensate stability, viscoelastic properties, and ageing behaviour can be systematically assessed. Understanding these mechanisms at the molecular level provides key insights into how specific mutations contribute to the dysregulation of phase separation, and paves the way for the rational design of targeted strategies to modulate biomolecular condensation therapeutically. In this context, machine learning algorithms can be instrumental in optimising sequence modifications that maximise the nucleation time of cross-*β*-sheet formation while preserving biological function. Moreover, engineering phase-separated organelles with finely tuned material properties holds great promise in synthetic biology and biotechnology, potentially enabling the development of novel drug delivery systems. Finally, predicting the molecular impact of sequence alterations—whether due to mutations or post-translational modifications—may offer critical insights into the relationship between condensate phase behaviour and disease, particularly in the case of FUS, where persistent stress granules have been linked to impaired cellular homeostasis and neurodegeneration.

## II. ACKNOWLEDGEMENTS

A. F. acknowledges funding from the Ramon y Cajal fellowship (RYC2021-030937-I) and Spanish National Grant (PID2022-136919NA-C33). I. S.-B. acknowledges funding from Derek Brewer scholarship of Emmanuel College and EPSRC Doctoral Training Programme studentship, number EP/T517847/1, Ramon y Cajal fellowship (awarded to J.R.E.). R.C.G, A.T., and I.S.B. acknowledge funding from the UK Research and Innovation (UKRI) Engineering and Physical Sciences Research Council (EPSRC) under the UK Government’s guarantee scheme (EP/Z002028/1), following successful evaluation by the ERC (Consolidator Grant awarded to R.C.G.) under the European Union’s Horizon Europe research and innovation programme. J. R. E. acknowledges funding from Emmanuel College, the University of Cambridge, and the Ramon y Cajal fellowship (RYC2021-030937-I). J. R. E. and F. G. also acknowledge the Spanish scientific plan and committee for research; project reference PID2022-136919NA-C33. E.P. acknowledges funding from an FPI scholarship associated to the Spanish scientific plan and committee for research: PID2022-136919NA-C33. This work has been performed using resources provided by the Cambridge Tier-2 system operated by the University of Cambridge Research Computing Service (http://www.hpc.cam.ac.uk) funded by EPSRC Tier-2 capital grant EP/P020259/1-CS170. This work has also been performed using resources provided by Archer2 (https://www.archer2.ac.uk/) funded by EPSRC Tier-2 capital grant EP/P020259/e829. The authors also thankfully acknowledge RES computational resources provided by Mare Nostrum 5 through the activities 2024-3-0001 and 2025-1-0009.

