## Supplementary Material for "Charged mutations in the FUS low-complexity domain modulate condensate ageing kinetics"

Eduardo Pedraza, Daniel Hoyos, Alejandro Feito, and Francisco Gámez
*Department of Physical Chemistry, Universidad Complutense de Madrid,*
*Av. Complutense s/n, Madrid 28040, Spain*

Ignacio Sanchez-Burgos
*Yusuf Hamied Department of Chemistry, University of Cambridge,*
*Lensfield Road, Cambridge CB2 1EW, UK*

Rosana Collepardo-Guevara
*Yusuf Hamied Department of Chemistry, University of Cambridge,*
*Lensfield Road, Cambridge CB2 1EW, UK and*
*Department of Genetics, University of Cambridge, Cambridge, UK*

Andrés R. Tejedor*
*Yusuf Hamied Department of Chemistry, University of Cambridge,*
*Lensfield Road, Cambridge CB2 1EW, UK and*
*Department of Physical Chemistry, Universidad Complutense de Madrid,*
*Av. Complutense s/n, Madrid 28040, Spain*

Jorge R. Espinosa†
*Department of Physical Chemistry, Universidad Complutense de Madrid,*
*Av. Complutense s/n, Madrid 28040, Spain and*
*Yusuf Hamied Department of Chemistry, University of Cambridge,*
*Lensfield Road, Cambridge CB2 1EW, UK*

(Dated: March 26, 2025)

### SI. THE MPIPI-RECHARGED MODEL

We use the Mpipi-Recharged model, a residue-level coarse-grained force field for both protein and RNA condensates [1]. In this model, each amino acid or nucleotide is represented by a single bead connected by harmonic bonds. Globular domains are treated as rigid bodies whose beads are fixed at the $C_\alpha$ of the corresponding Protein Data Bank (PDB). The potential energy is computed as the sum of pairwise bonded ($E_{\text{bonded}}$) and non-bonded ($E_{\text{non-bonded}}$) interactions as:

$$E = E_{\text{bonded}} + E_{\text{non-bonded}}. \tag{S1}$$

The intrinsically disordered regions (IDRs) are modelled as fully flexible polymers and these are connected to the globular domains. The bonded potential is written as:

$$E_{\text{bonded}}(r_{ij}) = \sum_{ij} k(r_{ij} - r_0)^2, \tag{S2}$$

where $r_{ij}$ is the distance between the connected beads, $r_0 = 3.81\,\text{Å}$ is the equilibrium bond length, and the spring constant $k = 9.6\,\text{kcal} \cdot \text{mol}^{-1} \cdot \text{Å}^{-2}$. The sum runs over all paired amino acids.

Non-bonded interactions consist of the sum of the hydrophobic interaction and electrostatic interaction. The hydrophobic interaction is given by the Wang–Frenkel (WF) potential [2] that accounts for short-ranged excluded-volume repulsion and long-ranged attraction. This potential is defined as

$$E_{\text{WF}}(r_{i,j}) = \sum_{ij} \epsilon_{ij}\alpha_{ij} \left[\left(\frac{\sigma_{ij}}{r_{ij}}\right)^{2\mu_{ij}} - 1\right] \left[\left(\frac{R_{ij}}{r_{ij}}\right)^{2\mu_{ij}} - 1\right]^{2\nu_{ij}}, \tag{S3}$$

where

$$\alpha_{ij} = 2\nu_{ij} \left(\frac{R_{ij}}{\sigma_{ij}}\right)^{2\mu_{ij}} \left\{ \frac{2\nu_{ij} + 1}{2\nu_{ij}\left[\left(\frac{R_{ij}}{\sigma_{ij}}\right)^{2\mu_{ij}} - 1\right]} \right\}^{2\nu_{ij}+1}. \tag{S4}$$

Here $\sigma_{ij}$ is the pair-of-beads diameter, defined from the individual diameter ($\sigma_i$ and $\sigma_j$) assuming the Lorentz-Berthelot mixing rules (i.e., $\sigma_{ij} = (\sigma_i + \sigma_j)/2$). $R_{ij} = 3\sigma_{ij}$ is the cut-off distance for the $ij$-th interaction. The interaction parameter $\epsilon_{ij}$ is defined for each specific amino acid pair based on our atomistic Potential of Mean Force calculations and

*
†

bioinformatics data [1]. The exponent $\mu_{ij}$ is set to 1 for all the pairs and $\nu_{ij}$ depends on the specific pair, ranging from 2 to 12 (see Ref. [1]). Notice that higher values of $\mu_{ij}$ lead to a steeper increase in the repulsive part of the potential.

The electrostatic interactions are described by the Yukawa potential [3] instead of the Debye–Hückel potential [4] originally employed in the Mpipi model. Avoiding the use of explicit charge values, this potential allows to modulate independently the strength of electrostatic interactions in a pair-specific basis. This potential is defined as

$$E_{\text{electrostatic}} = \sum_{ij} \frac{A_{ij}}{r_{ij}} \exp(-\kappa r_{ij}), \qquad r_{ij} < r_c, \tag{S5}$$

where $A_{ij}$ is the interaction parameter, $\kappa$ is the screening due to ions, and $r_{ij}$ is the distance between interacting pairs. The cut-off for the electrostatic interaction, $r_c$, is set to 3.5 nm. The screening parameter $\kappa$ is expressed in a explicit way as $\kappa = \sqrt{8\pi B c_s}$, where $c_s$ is the salt concentration—normally set to 150 mM of NaCl—and $B = e_0^2/4\pi k_B T \varepsilon_0 \varepsilon_r$ is the Bjerrum length. The relative dielectric constant $\varepsilon_r$ varies with temperature according to the empiric formula [5]:

$$\varepsilon_r(T) = \frac{5321}{T} + 233.760 - 0.9297T + 1.417 \cdot 10^{-3}T^2 - 8.292 \cdot 10^{-7}T^3, \tag{S6}$$

for $T$ in Kelvin. The pair-specific optimized values of $A_{ij}$ of the Yukawa potential can be found in Ref. [1]. In the Mpipi-Recharged model, such parameters indicate that the interaction between oppositely charged pairs is significantly stronger than those between identically charged pairs.

All simulations were carried out using the molecular dynamics LAMMPS software [6], version 2 of August of 2023.

### SII. FUS AND FUS-LCD SEQUENCES AND PDBS

#### FUS-LCD

MASNDYTQQATQSYGAYPTQPGQGYSQQSSQPYGQQSYSGYSQSTDTSGYGQSSYSSYGQSQNTG
YGTQSTPQGYGSTGGYGSSQSSQSSYGQQSSYPGYGQQPAPSSTSGSYGSSSQSSSYGQPQSGSYSQ
QPSYGGQQQSYGQQQSYNPPQGYGQQNQYNS

**FUS**

MASNDYTQQATQSYGAYPTQPGQGYSQQSSQPYGQQSYSGYSQSTDTSGYGQSSYSSYGQSQNTG
YGTQSTPQGYGSTGGYGSSQSSQSSYGQQSSYPGYGQQPAPSSTSGSYGSSSQSSSYGQPQSGSYSQ
QPSYGGQQQSYGQQQSYNPPQGYGQQNQYNSSSGGGGGGGGGNYGQDQSSMSSGGGSGGGYG
NQDQSGGGGSGGYGQQDRGGRGRGGSGGGGGGGGGYNRSSGGYEPRGRGGGRGGRGGMGGS
DRGGFNKFGGPRDQGSRHDSEQDNSDNNTIFVQGLGENVTIESVADYFKQIGIIKTNKKTGQPMIN
LYTDRETGKLKGEATVSFDDPPSAKAAIDWFDGKEFSGNPIKVSFATRRADFNRGGGNGRGGRGR
GGPMGRGGYGGGGSGGGGRGGFPSGGGGGGGQQRAGDWKCPNPTCENMNFSWRNECNQCKA
PKPDGPGGGPGGSHMGGNYGDDRRGGRGGYDRGGYRGRGGDRGGFRGGRGGGDRGGFGPGK
MDSRGEHRQDRRERPY

We have use two PDB codes to simulate the globular regions of FUS: residues from 285–371 (PDB code: 2LCW) and from 422–453 (PDB code: 6G99).

### SIII. CALCULATION OF THE PHASE DIAGRAM VIA DIRECT COEXISTENCE SIMULATIONS

The determination of the phase diagrams in the temperature-density plane was performed using the Direct Coexistence method [7, 8]. The proteins are placed in a prismatic elongated box to simulate both the high-density and low-density phases separated by an interface. The long side of the box is perpendicular to the interfaces. Since determination of the optimal box dimensions is key to minimising finite size effects while ensuring computer efficiency, we provide the following guidelines

1. A minimum of 48 protein replicas for full FUS systems and a minimum of 80 protein replicas for FUS-LCD systems per box were used.

2. To avoid self-interactions through periodic boundary conditions we enforce the short sides of the box to be larger than at least twice the radius of gyration of the protein in the system.

3. The long side of the box should keep the total density of the system at approximately $\sim 0.1\,\mathrm{g{\cdot}cm^{-3}}$.

Simulations are carried out in the canonical (NVT) ensemble using a Nosé-Hover thermostat [9] for the rigid bodies (representing the globular domains) integrated in the RIGID package, and a Langevin thermostat [10] for the particles in flexible regions, both with a relaxation time of 5 ps. The timestep for the Verlet integration of the equations of motion is 10 fs. After equilibration has reached, we run the simulation of 1-2 $\mu$s depending on the specific system. If two different phases are detectable—i.e., a high density phase and a low density phase—, the densities are calculated.

The critical density ($\rho_c$) and temperature ($T_c$) of the phase diagrams are evaluated by means of the law of rectilinear diameters and critical exponents [11, 12]:

$$\frac{\rho_l(T) + \rho_d(T)}{2} = \rho_c + s_2(T_c - T), \tag{S7}$$

and

$$(\rho_l(T) - \rho_d(T))^\beta = d\left(1 - \frac{T}{T_c}\right), \tag{S8}$$

where the critical exponent $\beta = 3.06$, $\rho_d$ and $\rho_l$ are the coexisting densities of the diluted and condensed phases, respectively, and $d$ and $s_2$ are fitting parameters.

### SIV. DYNAMIC ALGORITHM FOR CONDENSATE AGEING SIMULATIONS

We use a dynamic ageing algorithm [13, 14] to enable 'effective' disorder-to-order transitions by forming inter-protein $\beta$-sheets between the low-complexity aromatic-rich kinked segments (LARKS, i.e., the segments $_{37}$SYSGYS$_{42}$, $_{54}$SYSSYGQS$_{61}$, and $_{77}$STGGYG$_{82}$) of FUS and FUS-LCD and monitor the kinetics of the process depending on the local environment. The algorithm checks every $10^2$ simulation time steps the local environments of the LARKS and, when five LARKS within the coordination cut-off distance ($r_{cut}$, see Table S1), the dynamic algorithm changes the force field parameters ($\epsilon_{ij}$,$\sigma_{ij}$) of the bead contained in the LARKS to those ($\epsilon_{ij,ordered}$,$\sigma_{ij,ordered}$) given in Table S1. The parameters correspond to inter-protein structured $\beta$-sheets as obtained from atomistic Potential of Mean Force simulations (see Refs. [13–15]). The formed inter-protein $\beta$-sheet can grow if another LARKS is recruited within the $r_{cut,cross}$ distance (see Table S1). Similarly, an inter-protein $\beta$-sheet is reverted if one of the structured LARKS separates from the crossed-$\beta$-sheet motif a distance ($r_{cut,rev}$ away, see Table S1). Additionally, the algorithm increases the local stiffness to mimic the structured $\beta$-sheet by introducing a harmonic angular potential given by

$$E_{\text{angle}} = \sum_{\text{angles}} k_{ang}(\theta - \theta_0)^2, \tag{S9}$$

where we set $\theta_0 = 180°$ and $k_{ang}$=5 kcal mol$^{-1}$ rad$^{-2}$ for the structured LARKS of an inter-protein $\beta$-sheet.

These simulation were performed using the bond/react fix available in the REACTION package [16] of LAMMPS version 2 Aug 2023, which allows us to change the topology and identity of the selected residues (i.e., LARKS within a LARKS high-density fluctuation) in a time- and local-dependent way. We run the dynamic algorithm in the isothermal-isobaric (NpT) ensemble for full FUS systems and in NVT for FUS-LCD systems using bulk conditions, with a relaxation time of 5 ps for the Langevin thermostat and the Nosé Hoover barostat, and a time step of 10 fs.

| LARKS of FUS-LCD | $_{37}$SYSGYS$_{42}$ | $_{54}$SYSSYGQS$_{61}$ | $_{77}$STGGYG$_{82}$ |
|---|---|---|---|
| Central amino acid | $S_{39}$ | $S_{57}$ | $G_{79}$ |
| $m_{ordered}$ / (g mol$^{-1}$) | 107.44 | 107.48 | 86.92 |
| $\epsilon_{ij,ordered}$ / (kcal mol$^{-1}$) | 1.551 | 3.113 | 0.892 |
| $\sigma_{ij,ordered}$ / Å | 5.733 | 5.761 | 5.353 |
| $r_{cut}$ / Å | 14 | 14 | 14 |
| $r_{cut,cross}$ / Å | 13 | 13 | 13 |
| $r_{cut,rev}$ / Å | - | - | 20 |
| Coordination number | 5 | 5 | 5 |

**TABLE S1:** Parameters employed for residues belonging to structured inter-peptide $\beta$-sheet motifs in Mpipi-Recharged simulations for the LARKS in FUS-LCD, including the mass $m_{ordered}$, the interaction $\epsilon_{ij,ordered}$ and the steric radius $\sigma_{ij,ordered}$ of the structured LARKS, the cut off distances ($r_{cut}$, $r_{cut,cross}$, $r_{cut,rev}$), and the number of LARKS necessary to trigger the algorithm (coordination number).

For the sake of computational feasibility, the cut-off distances for FUS condensates are longer so that the disorder-to-order transitions in full FUS present a similar timescale to that in FUS-LCD (see Table S2). The number of LARKS necessary to form $\beta$-sheets is reduced to 4 LARKS within the coordination sphere. The results are presented in Figure S1.

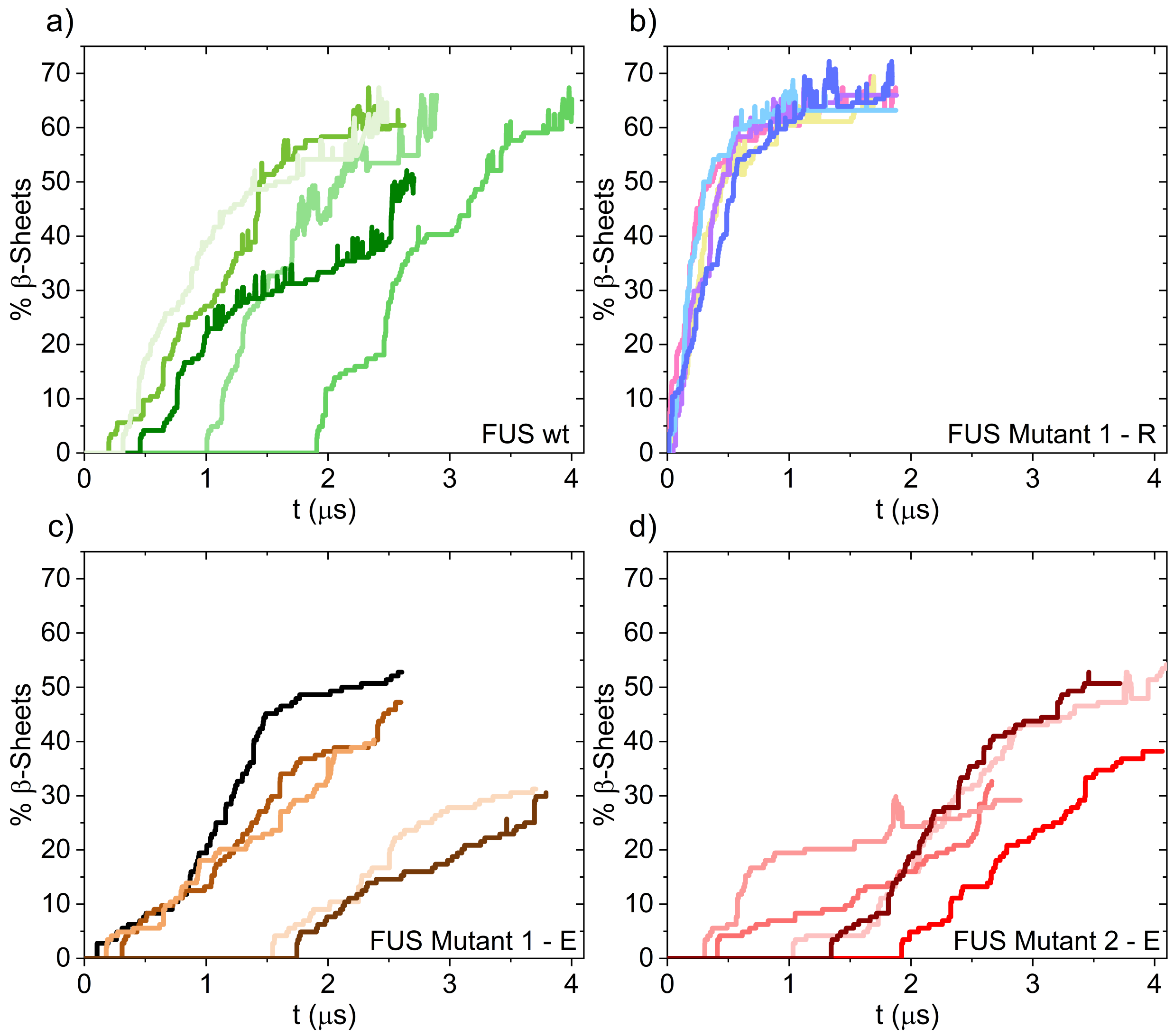


**FIG. S1:** The percentage of interprotein $\beta$-sheet transitions as a function of time is presented for a condensate of full FUS wild-type (a) and for three full FUS mutants condensates: In panel (b), Mutant 1 with arginine in panel (c), Mutant 1 with glutamic acid; and in panel (d), Mutant 2 with glutamic acid.

| LARKS of FUS-LCD | $_{37}$SYSGYS$_{42}$ | $_{54}$SYSSYGQS$_{61}$ | $_{77}$STGGYG$_{82}$ |
|---|---|---|---|
| Central amino acid | $S_{39}$ | $S_{57}$ | $G_{79}$ |
| $m_{ordered}$ / (g mol$^{-1}$) | 107.44 | 107.48 | 86.92 |
| $\epsilon_{ij,ordered}$ / (kcal mol$^{-1}$) | 1.551 | 3.113 | 0.892 |
| $\sigma_{ij,ordered}$ / Å | 5.733 | 5.761 | 5.353 |
| $r_{cut}$ / Å | 12 | 12 | 12 |
| $r_{cut,cross}$ / Å | 12.5 | 12.5 | 12.5 |
| $r_{cut,rev}$ / Å | - | - | 20 |
| coordination number | 4 | 4 | 4 |

**TABLE S2:** Parameters employed for residues belonging to structured inter-peptide $\beta$-sheet motifs in Mpipi-Recharged simulations for the LARKS in FUS, including the mass $m_{ordered}$, the interaction $\epsilon_{ij,ordered}$ and the steric radius $\sigma_{ij,ordered}$ of the structured LARKS, the cut off distances ($r_{cut}$, $r_{cut,cross}$, $r_{cut,rev}$), and the number of LARKS necessary to trigger the algorithm (coordination number).

### SV. ESTIMATION OF NUCLEATION TIMES

As observed in the main text, some condensates did not rigidify during the course of the 3 $\mu s$ simulations. To evaluate the nucleation times at which $\beta$-sheet formation would begin, we assumed that the rigidification process follows first-order equation kinetics given by

$$\ln\left(\frac{N(t)}{N(0)}\right) = -kt, \tag{S10}$$

where $N(t)$ is the number of liquid condensates at time $t$ and $k$ is the rate constant that characterizes the rigidification rate of the condensates. In our work $N(0) = 5$ (we have 5 condensates in each set of FUS-LCD mutations) and, except in the case of the mutant 2 - E, at least two condensates rigidified for each protein. Thus, $N(t)$ can be 1, 2, or 3. Therefore, we can plot the values of $\ln\left(\frac{N(t)}{N(0)}\right)$ vs $t$, fit these data to a regression line and estimate, through extrapolation, the nucleation times at which condensates that remain liquid after 3 $\mu$s would begin to form interprotein $\beta$-sheets. For the validation of this approach (Fig. S2.a), the nucleation times of the mutant 1-R (in which all condensates formed $\beta$-sheets, solid points in the Fig. S2) were used. To do this, the nucleation time of two condensates (hollow points in Fig. S2.a) was determined from the three lowest nucleation times. Finally, the average nucleation times were determined, both real (using the nucleation times measured in the simulation, all solid points in the Fig. S2.a), $\tau_{real}$, and estimated (using the nucleation times used for fitting and the two estimates derived from them), $\tau_{estimated}$. This determination yielded similar values for both $\tau_{real}$ and $\tau_{estimated}$, thereby validating the approximation. As example, the estimation for mutant 4-E is included in Fig. S2.b.

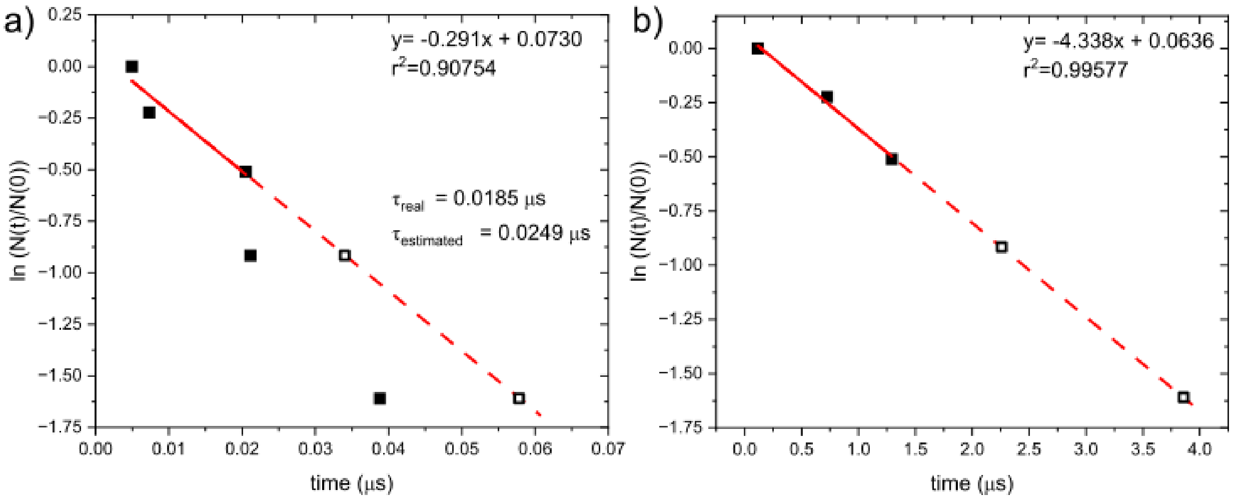


**FIG. S2:** a) Validation of the kinetic approximation for Mutant 1-R. b) Determination of nucleation times for the liquid condensates after 3 $\mu$s of simulation for Mutant 4-E.

### SVI. CALCULATION OF THE VISCOSITY

In an isotropic system, the Green-Kubo relation allow to obtain the viscosity, $\eta$, by integrating the shear relaxation modulus $G(t)$, i.e., $\eta = \int_0^\infty G(t)dt$. $G(t)$ can be calculated after evaluating the components of the pressure tensor ($\sigma_{\alpha\beta}$) as arising from the virial theorem, as follows [17]:

$$\begin{aligned} G(t) = &\frac{V}{5k_BT}\left[\langle\sigma_{xy}(0)\sigma_{xy}(t)\rangle + \langle\sigma_{xz}(0)\sigma_{xz}(t)\rangle + \langle\sigma_{yz}(0)\sigma_{yz}(t)\rangle\right] \\ &+ \frac{V}{30k_BT}\left[\langle N_{xy}(0)N_{xy}(t)\rangle + \langle N_{xz}(0)N_{xz}(t)\rangle + \langle N_{yz}(0)N_{yz}(t)\rangle\right], \end{aligned} \tag{S11}$$

where $N_{\alpha\beta} = \sigma_{\alpha\alpha} - \sigma_{\beta\beta}$ is the normal stress difference. Finally, $\eta$ is obtained following a two-step procedure. First, we numerically integrate the first part of the decay of $G(t)$ up to $t_0$, thus obtaining the intramolecular contribution to the viscosity ($\eta_0$). Second, in the long time limit, the slower relaxation is integrated by fitting $G(t)$ to the sum of $m$=4–6 Maxwell modes, depending on the span and noise of $G(t)$. The discrete Maxwell modes have the exponential form $G_i \exp(-t/\tau_i)$ and are equidistant in logarithmic time [18]. The fit is carried out with the help of the open-source RepTate software [19]. Finally, viscosity is obtained by adding the two terms:

$$\eta = \eta_0 + \sum_{i=0}^{m} \tau_i G_i \exp\Big(-t/\tau_i\Big) \tag{S12}$$

where $\tau_i$ and $G_i$ are the fitting parameters of $G(t)$ written as the sum of Maxwell modes. Typically, simulations of 3-5 $\mu$s are needed to observe a clear terminal decay in $G(t)$, characteristic of liquid phases. Storage ($G'$) and loss ($G''$) moduli can be directly obtained as the Fourier transform of $G(t)$ also using RepTate [19].

The components of the pressure tensor were evaluated in bulk conditions at the critical densities in the NVT ensemble employing a Langevin thermostat with a relaxation time of 5 ps and a time step of 10 fs. The configurations were generated by placing 100 protein replicas for FUS-LCD systems and 48 protein replicas for full FUS systems in a cubic box.

### SVII. ANALYSIS OF THE NETWORK CONNECTIVITY BY A PRIMITIVE PATH ANALYSIS

We analyse the $\beta$-sheet network connectivity of the aged condensates using the primitive path analysis (PPA) method [20] modified in Tejedor *et al.* [14]. The PPA algorithm minimises the contour length of all chains in the condensate, while keeping the terminal residues of the monomers fixed, thus keeping the network topology by impeding chains to cross each other. First, we perform a short NpT simulation to allow the number of cross-$\beta$-sheet transitions to reach the equilibrium density. From this configuration, the positions of the intermolecular $\beta$-sheet motifs are frozen, and then, we performed an energy minimisation of $10^5$ iterations with a tolerance of $10^{-6}$ for both energy and total force. In this step, the bond length is fixed to zero and the excluded volume interactions are removed so that the contour length of the strands between the transitioned structured LARKS is minimised while preserving the underlying network connectivity.

### SVIII. CALCULATION AND ANALYSIS OF CONTACT MAPS

The calculation of intermolecular contact maps within protein condensates is calculated from NVT trajectories. Contacts were determined in all systems at 290 K. Typically, molecular contacts are identified based on a distance criterion, with the assumption that the relative frequency of contact map occurrences (rather than absolute frequency) remains generally unaffected by the selected cut-off distance used in calculations, provided the cut-off values are reasonable. However, to accurately capture the most relevant and common residue-residue contact pairs that facilitate LLPS, it is highly recommended to account for the specific parameterisation of each amino acid in terms of excluded volume and minimum potential energy interaction distance. Hence, we adopted a sequence-dependent cut-off distance equivalent to $1.2\sigma_{ij}$, where $\sigma_{ij}$ represents the mean excluded volume of the respective i-th and j-th amino acid [21]. Given that the minimum of the potential used is approximately $\sqrt[6]{2}\sigma_{ij} \approx 1.122\sigma_{ij}$, we set the cut-off distance slightly beyond this point, at $1.2\sigma_{ij}$, to ensure significant binding. By implementing this innovative sequence-dependent cut-off scheme for each amino acid pair interaction, we can effectively filter out adjacent contacts that may coincide with actual interacting amino acids along the sequence, thus enhancing our ability to accurately identify the amino acids that positively contribute to stabilising condensates [21].

The resulting intermolecular contact maps show the total contact frequency between the amino acids of different proteins within a condensate. To analyse in detail the variation in contact between the LARKS of the proteins, the following formula has been used

$$\% \text{ Contact Variation} = \frac{M - W}{W} \cdot 100, \tag{S13}$$

where M represents the sum of intermolecular contacts of the LARKS of interest in the mutated protein, and W represents the sum of intermolecular contacts of the same LARKS in the protein in its wild-type form. In the intermolecular contact maps of full FUS condensates, since the total number of contacts may vary due to small changes in system density, M and W must be normalised by the total sum of contacts in the corresponding intermolecular contact map. In the FUS-LCD maps, this normalization is not necessary.

This formula can be used not only to analyse the percentage variation in contact between LARKS of the mutated protein relative to the wild-type, but it can also be applied to analyse variations in any part of the protein sequence. The following tables summarise the percentage variation in interaction between LARKS relative to the wild-type, in the different mutants of FUS-LCD and full FUS.

| | $^{37}$SYSGYS$^{42}$ | $^{54}$SYSSYGQS$^{61}$ | $^{77}$STGGYG$^{82}$ |
|---|---|---|---|
| Mutant 1 - E | -33.76 | -41.81 | -25.19 |
| Mutant 2 - E | -51.03 | -37.37 | -14.80 |
| Mutant 1 - R | 121.11 | 106.25 | 66.00 |
| Mutant 1 - A | -11.49 | -13.21 | -15.54 |
| Mutant 1 - K | -22.40 | -40.17 | -26.08 |
| Mutant 1 - D | -39.15 | -44.87 | -17.76 |
| Mutant 3 - E | -20.71 | -23.21 | -16.25 |
| Mutant 4 - E | -32.48 | -25.40 | -13.22 |

**TABLE S3:** % Contact variation in LARKS relative to the wild-type for FUS-LCD mutants.

| | $^{37}$SYSGYS$^{42}$ | $^{54}$SYSSYGQS$^{61}$ | $^{77}$STGGYG$^{82}$ |
|---|---|---|---|
| Mutant 1 - E | -0.81 | 0.06 | –25.10 |
| Mutant 2 - E | -11.41 | -14.59 | -7.80 |
| Mutant 1 - R | 141.10 | 126.95 | 69.17 |

**TABLE S4:** % Contact variation in LARKS relative to the wild-type for full FUS mutants.

| | LCD-LCD | LCD-RGG1 | LCD-RRM | LCD-RGG2 | LCD-ZF | LCD-RGG3 | LCD-Residue |
|---|---|---|---|---|---|---|---|
| Mutant 1 - E | -2.79 | 5.00 | -2.53 | 14.87 | 3.98 | 1.53 | 3.80 |
| Mutant 2 - E | -5.49 | 6.02 | -2.04 | 13.04 | 1.86 | 0.86 | 3.45 |
| Mutant 1 - R | 103.63 | -20.27 | -2.69 | -36.86 | -17.84 | -12.43 | -17.55 |

**TABLE S5:** % Contact variation in LARKS relative to the wild-type for full FUS protein domains.

### SIX. CONTACT MAPS OF FUS-LCD MUTANTS

The intermolecular contact maps of these FUS-LCD mutants were calculated from simulations in the NVT ensemble of 3 $\mu$s, as detailed in section SVIII. All maps are expressed in residue-residue % contact frequency.

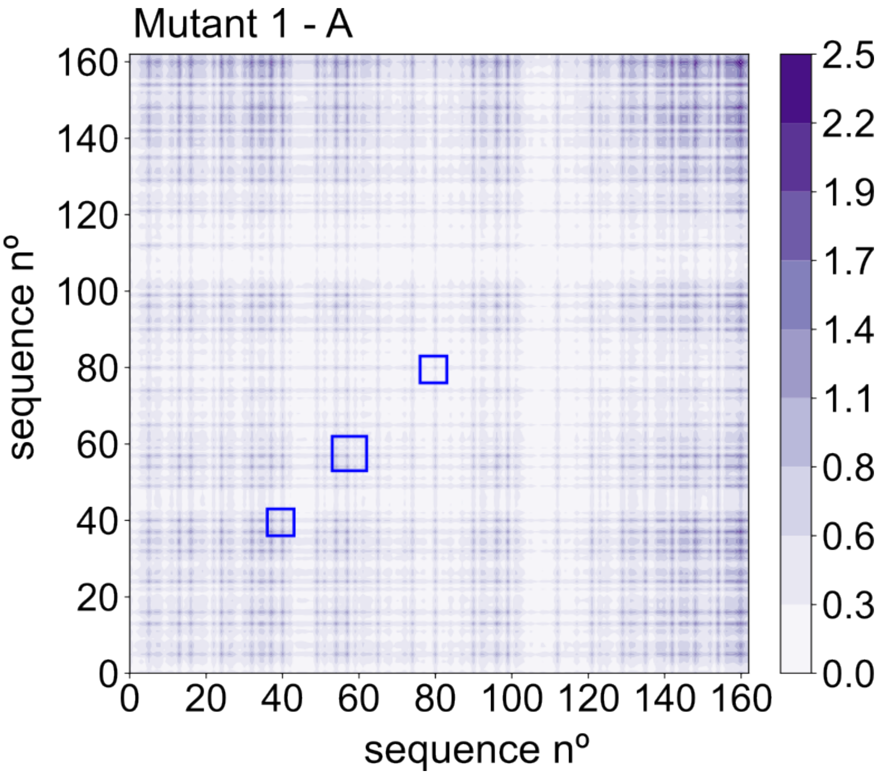


**FIG. S3:** Intermolecular contact map probability prior ageing for FUS-LCD Mutant 1-A.

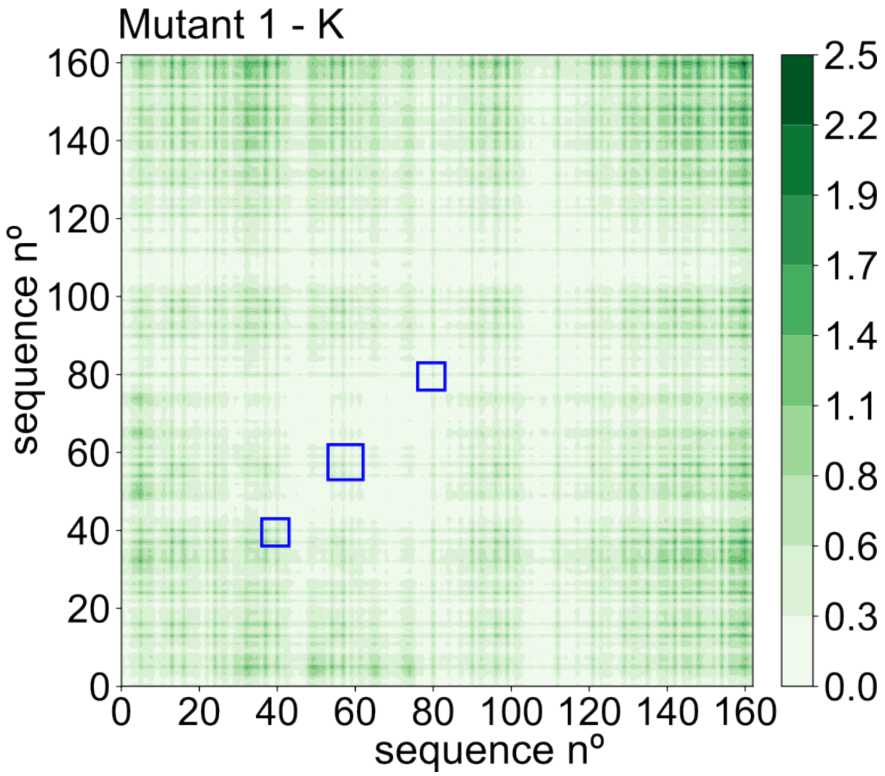


**FIG. S4:** Intermolecular contact map probability prior ageing for FUS-LCD Mutant 1-K.

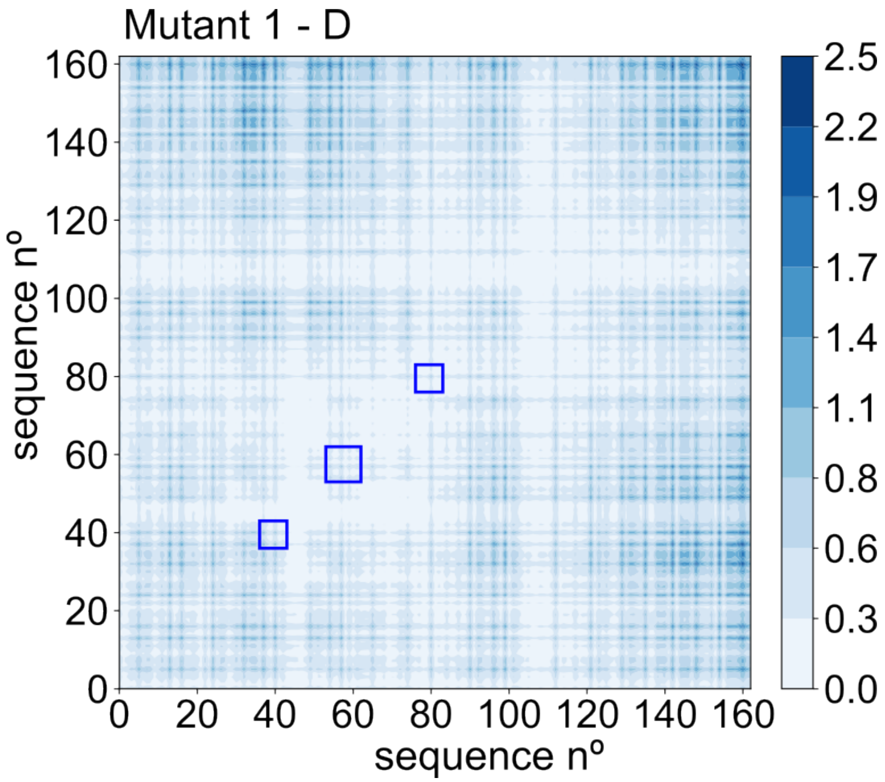


**FIG. S5:** Intermolecular contact map probability prior ageing for FUS-LCD Mutant 1-D.

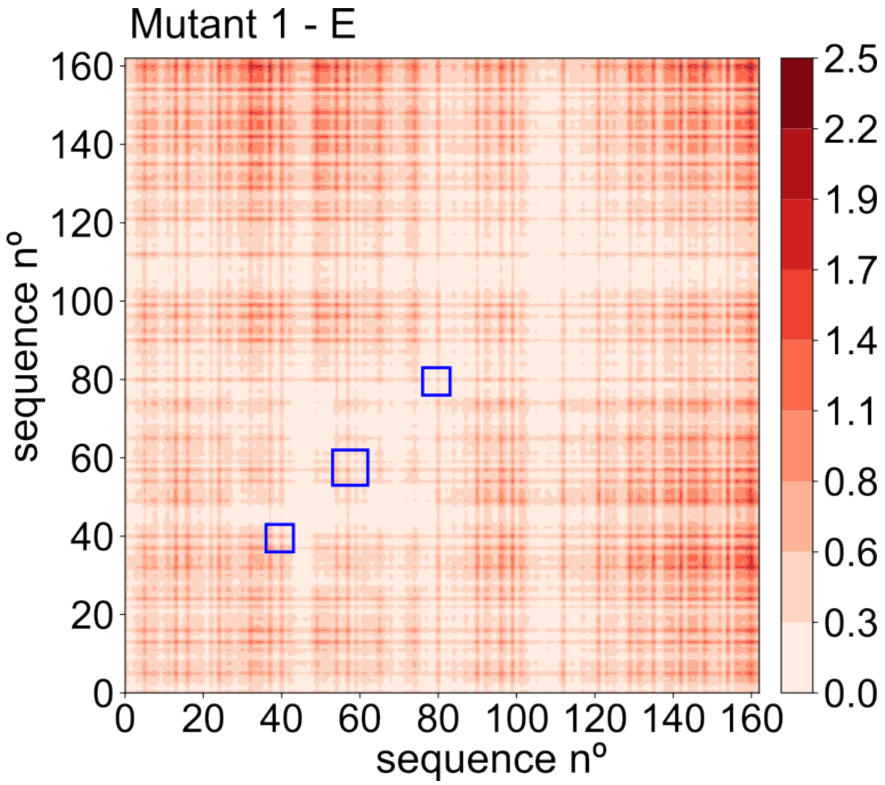


**FIG. S6:** Intermolecular contact map probability prior ageing for FUS-LCD Mutant 1-E.

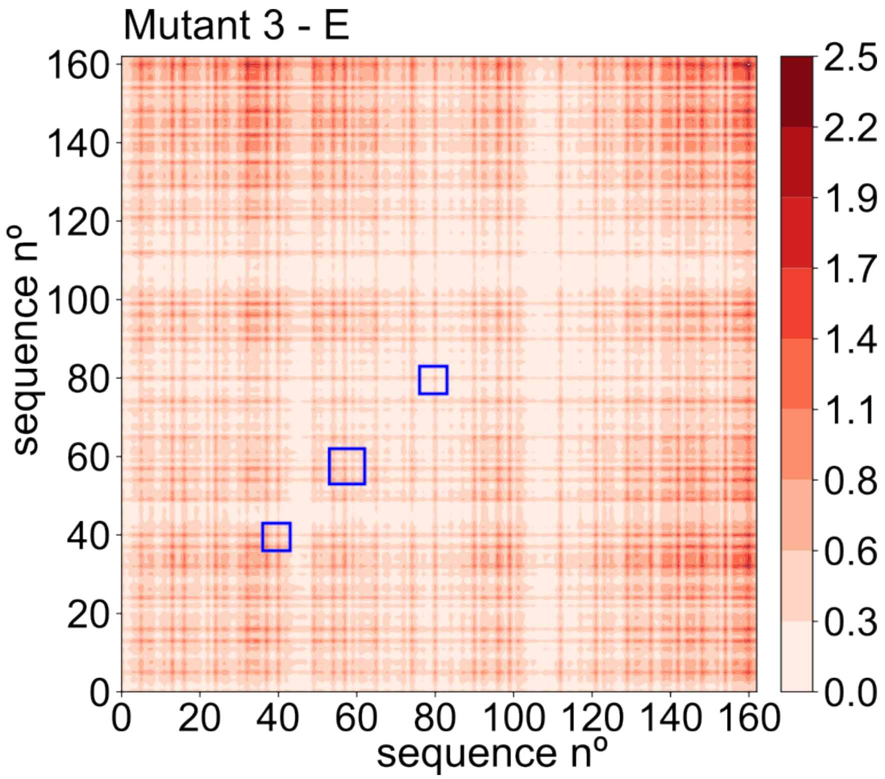


**FIG. S7:** Intermolecular contact map probability prior ageing for FUS-LCD Mutant 3-E.

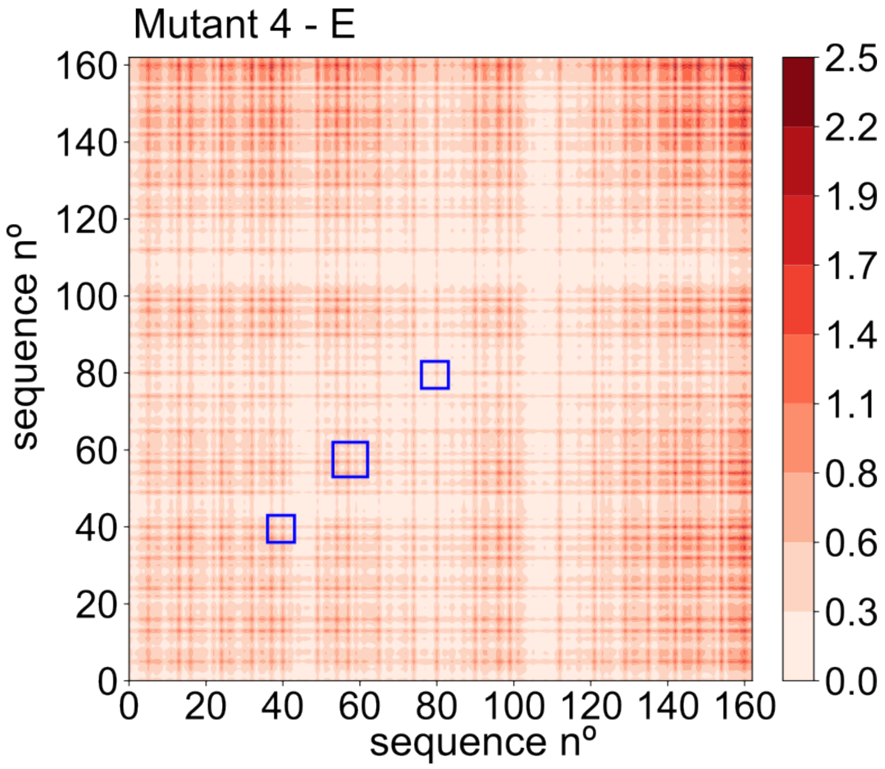


**FIG. S8:** Intermolecular contact map probability prior ageing for FUS-LCD Mutant 4-E.

[12] J. S. Rowlinson and B. Widom, *Molecular theory of capillarity.* Courier Corporation, 2013.
